# Transient EZH2 suppression by Tazemetostat during *in vitro* expansion maintains T cell stemness and improves adoptive T cell therapy

**DOI:** 10.1101/2023.02.07.527459

**Authors:** Yingqin Hou, Jaroslav Zak, Yujie Shi, Isaraphorn Pratumchai, Brandon Dinner, Wenjian Wang, Ke Qin, Evan Weber, John R. Teijaro, Peng Wu

## Abstract

The histone methyltransferase enhancer of zeste homolog 2 (EZH2)-mediated epigenetic regulation of T cell differentiation in acute infection has been extensively investigated. However, the role of EZH2 in T cell exhaustion remains under-explored. Here, using *in vitro* exhaustion models, we demonstrated that transient inhibition of EZH2 in T cells before the phenotypic onset of exhaustion with a clinically approved inhibitor, Tazemetastat, delayed their dysfunctional progression and maintained T cell stemness and polyfunctionality while having no negative impact on cell proliferation. Tazemetestat induced T cell epigenetic reprogramming and increased the expression of the self-renewing T cell transcription factor TCF1 by reducing its promoter H3K27 methylation preferentially in rapidly dividing T cells. In a murine melanoma model, T cells pre-treated with tazemetastat exhibited a superior response to anti-PD-1 blockade therapy after adoptive transfer. Collectively, these data unveil the potential of transient epigenetic reprogramming as a potential intervention to be combined with checkpoint blockade for immune therapy.

## Introduction

Adoptive T cell therapy (ACT)-based cancer immunotherapies, which is the infusion of autologous or allogeneic T cells including tumor-infiltrating lymphocytes (TILs), T cell receptor T cells (TCR-T), and chimeric antigen receptor T cells (CAR-T) into patients after *ex vivo* expansion, have induced remarkable clinical responses in patients with hematopoietic malignancies.^[1]^ However, long-lasting responses are limited to a subset of individuals. Furthermore, it has proven difficult to achieve significant and sufficient antitumor activities by employing these approaches to treat solid tumors.^[2]^ The occurring of exhaustion and functional impairment of T cells used for ACT is one of the major roadblocks that limit the successful application of ACT for treatment of solid tumors.

During chronic viral infections and cancer, persistent antigen exposure drives T cell exhaustion, whose manifestation includes progressive and hierarchical loss of effector functions, sustained upregulation and co-expression of multiple inhibitory receptors (e.g. PD-1, LAG3, Tim-3, 2B4, TIGIT, CD39),^[3, 4]^ altered expression and use of key transcription factors (e.g. TCF-1, TOX, NFAT1, IRF4, BATF), ^[5-8]^metabolic defects (e.g. impaired OXPHOS, high ROS production),^[9, 10]^ and a failure to transition to quiescence and acquire antigen-independent memory T cell homeostatic responsiveness. Likewise, during *ex vivo* expansion regimens used to grow a small fraction of reactive T cells into large numbers for patient infusion, T cells are often driven into terminal differentiation and exhaustion. Consequently, proliferation potential after adoptive transfer is very limited or absent. Recent studies revealed that two distinct populations of exhausted CD8^+^ T cells exist: progenitor (PD-1^+^Tim-3^−^TCF-1^+^) and terminally (PD-1^+^Tim-3^+^TCF-1^−^) exhausted T cells.^[11-15]^ Progenitor exhausted T cells exhibit stem cell-like properties and give rise to terminally exhausted CD8^+^ T cells (T_term-ex_) that possess cytotoxic functions. During this differentiation process, the transcription factor TCF-1 (T cell factor 1 encoded by *Tcf7*), playing a central role in the transcriptional network that drives a stem-like versus terminally exhausted T cell-fate decision, ^[5, 16]^ is gradually lost. Progenitor exhausted T cells possess self-renewable capabilities and can respond to anti-PD-1 therapy, but terminally exhausted T cells lack such properties.

T cell activation and differentiation are accompanied by epigenetic changes that switch genes on or off and determine which proteins are transcribed.^[17, 18]^ Such changes in turn play pivotal roles in the determination of T cell fate and function. DNA methylation and histone post-translational modifications represent two major epigenetic mechanisms. Enhancer of zeste homolog 2 (EZH2), the functional enzymatic component of the Polycomb Repressive Complex 2 (PRC2), is a histone-lysine N-methyltransferase enzyme encoded by *Ezh2* gene.^[19, 20]^ It mediates gene repression by catalyzing trimethylation of histone H3 at Lys27 (H3K27me3).

Previous studies in a murine melanoma model revealed that CD8^+^ T cells lacking EZH2 had reduced cytokine production and poor tumor control.^[21]^ Similarly, using an acute viral infection model, Kaech et al. uncovered that *Ezh2* deficiency led to a compromised CD8^+^ effector program without jeopardizing memory cell formation.^[22]^ Analyzing *Ezh2* targets whose expression decreased during the differentiation of effector cells but not during the differentiation of memory cells revealed that many genes encode products that have been linked to the memory formation but not to the effector function.^[22, 23]^ These Ezh2 targets included genes encoding memory-associated transcription factors, such as *Tcf7, Bach2*, and *Eomes*; molecules that mediate TGF-signaling, e.g., *Smad2*, whose product has been linked to CD8+ T cell fate ‘decision’;^[24-26]^ factors that control T cell survival and homing, e.g., *Bcl2* and *Klf2*;^[27, 28]^ as well as *Opal1*, which encodes a regulator of mitochondrial fusion with a critical role in differentiating memory CD8^+^ T lymphocytes.^[29]^ Although the role of EZH2 in the establishment of the epigenetic state of exhausted T cells remains to be explored, many genes highly expressed in progenitor exhausted T cells overlap with the aforementioned EZH2 targets that are linked to the differentiation of memory cells but not to the differentiation of effector cells.^[14]^ These observations suggest that transient inhibition of EZH2 in the early stage of T cell differentiation toward exhaustion may facilitate the generation of progenitor exhausted T cells with “stem-like” properties that are highly desirable for cancer immunotherapy in the settings of immune checkpoint blockade and ACT.

Tazemetostat (Taz) is a S-adenosyl methionine (SAM) competitive inhibitor of EZH2 approved by FDA in 2020 for the treatment of metastatic or locally advanced epithelioid sarcoma not eligible for complete resection and relapsed/refractory follicular lymphoma.^[30, 31]^ We hypothesize that culturing CD8^+^ T cells with Taz during *ex vivo* expansion would inhibit the deposit of H3K27Me3 at memory associated loci, alleviate exhaustion, and produce T cells with stem-like or progenitor-exhausted properties. Treating T cells with Taz confines inhibition of EZH2 to *ex vivo* expansion, thus avoiding any long-term functional EZH2 loss after adoptive transfer. Upon adoptive transfer, in the absence of Taz, EZH2 activity would be recovered, enabling the transferred T cells to differentiate into cells with effector-like functions for better tumor control.

## Results

### EZH2 expression is associated with T cell exhaustion and proliferation

Exhausted T cells harbor distinct epigenetic features distinct from that of effector and memory T cells which are minimally remodeled after checkpoint blockade. Therefore, as the first step to investigate the role of EZH2 in establishing T cell exhaustion, we searched publicly accessible sequencing data to explore whether the expression of EZH2 is associated with distinct immune states such as exhaustion. Interestingly, whereas the expression of EZH2 in virus-specific CD8 T cells during acute infection peaked and then returned to normal levels after virus clearance, *Ezh2* levels persisted at above-naïve levels in T cells during persistent LCMV Clone 13 infection (**Figure 1A0**). This effect could be partly explained by changes in cell cycle, differentiation, or both. We examined public single-cell RNA-seq datasets to further resolve the factors associated with *Ezh2* expression. Interestingly, a broad panel of human and animal studies revealed that EZH2 expression was not associated with specific T cell states, but rather with actively cycling cells (**Figure 1A-F**). For example, all 11 non-cycling T cell subsets examined in a human breast cancer dataset showed comparably low levels of *Ezh*2, whereas cycling T cells and other cycling cell types showed high levels of *Ezh2* (**Figure 1C**). Accordingly, there was no association of *Ezh2* expression with T cell exhaustion or activation markers such as *PDCD1* or *CD69*, but a very strong association with mitotic genes such as *Mki67* or *CENPA* (**Figure 1A-F**). The statistically significant and cross-cell-type association of expression *Ezh2* with mitosis was confirmed in bulk tumor RNA-seq datasets from multiple TCGA cohorts (Figure 1G-H). Moreover, mouse *Ezh2* showed the highest expression in thymocytes among mouse T cell samples, and the relative levels of *Ezh2* compared to *Mki67* were the highest in double positive thymocytes (Figure 1I). These observations validate earlier reports of mitosis-associated *Ezh2* expression,^[32]^ and suggest that high *Ezh2* levels mark T cell subsets with the highest percentage of actively dividing cells.

**Figure 1:**
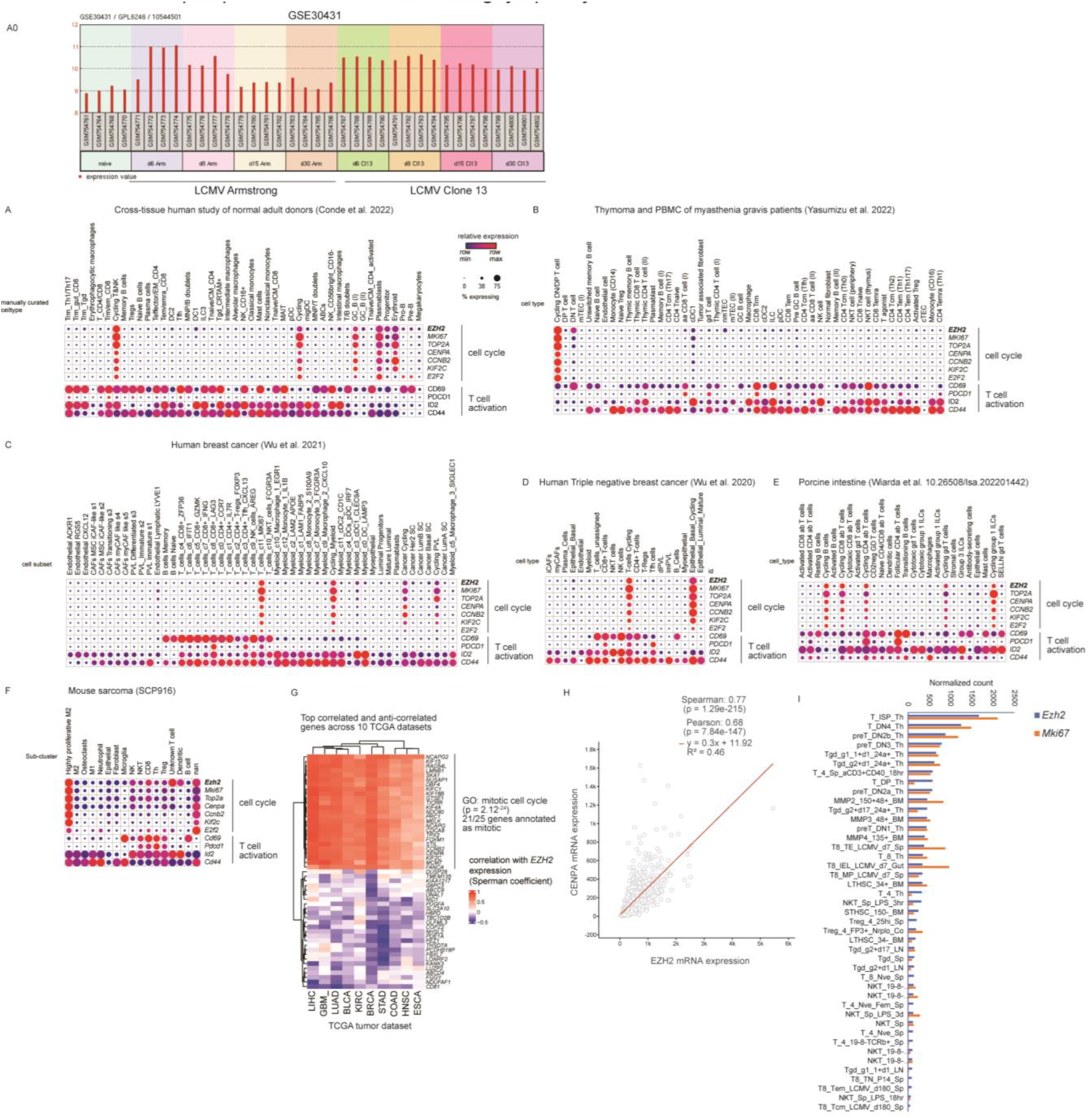
(**A-F**) Expression levels of *EZH2*, selected mitotic genes and T cell activation in publicly available single-cell RNA-seq datasets; size of circle indicates percentage of cells expressing transcript, color indicates level of expression; (**G**) Genes most highly correlated (positively and negatively) with *EZH2* expression in human tumor bulk RNA-seq datasets, ranked by mean Spearman rank correlation coefficient, top 25 genes plotted; (**H**) expression of mitotic gene *CENPA* vs *EZH2* in breast cancer, batch normalized values shown; (**I**) normalized expression of *Ezh2* and *Mki67* in T cell bulk RNA-seq Immgen samples; BLCA, bladder carcinoma; BRCA, invasive breast carcinoma; COAD, colorectal adenocarcinoma; ESCA, esophageal carcinoma; GBM, glioblastoma; HNSC, head and neck squamous cell carcinoma; LIHC, liver hepatocellular carcinoma; LUAD, lung adenocarcinoma; KIRC, kidney renal clear cell carcinoma; STAD, stomach adenocarcinoma; DN, double negative; Sp, spleen; Th, thymic cells; T8, CD8 T cells; T4, CD4 T cells; BM, bone marrow.

### Transient inhibition of EZH2 with Taz promotes progenitor-like features of T cells without compromising cell expansion

As described above, actively dividing T cells including those destined for exhaustion have high levels of EZH2 expression and EZH2 mediates the deposition of the suppressive H3K27Me3 mark on memory and stemness related gene. Therefore, we tested if transient inhibition of the function of EZH2 by treating T cells in the early stage of differentiation toward exhaustion with Taz may halt or slow their exhaustion progression and promote the generation of less-differentiated T cells with stem-like features. Toward this goal, we developed an *ex vivo* T cell exhaustion model using OT-I T cells that express a transgenic TCR specific for the SIINFEKL peptide (OVA_257–264_) of chicken ovalbumin presented on the MHC I molecule H2-K^b^ (**Figure 2A**). In this assay, OT-I splenocytes were primed with a high concentration (500 nM) of OVA_257–264_ for 3 days, followed by *ex vivo* expansion, which resulted in pronounced T cell exhaustion by day 7, featuring the up-regulation of inhibitory receptors PD-1, LAG3, Tim-3, 2B4, and CD39, decreased cytokine production, and slower T cell proliferation as compared with OT-I cells activated with a low concentration (1 nM) of OVA_257–264_ that possess an effector memory phenotype (**Figure 2A** and **Figure S1**). Upon restimulation *in vitro* on day 7, OT-I T cells primed with the high concentration of OVA_257–264_ underwent pronounced apoptosis, and dramatically reduced the capabilities to trigger the killing of 4MOSC1-SIINFEKL cells in comparison to their counterparts primed with 1 nM of OVA_257–264_ (**Figure S1 G-J**). Worthy of note, in the absence of Taz during expansion, by day 5, 29.0% OT-I CD8^+^ T cells were terminally exhausted (Tim3^+^TCF-1^−^) and only 20.4% exhibited progenitor exhausted phenotype (Tim3^−^TCF-1^+^) (**Figure 2B**). By contrast, when Taz was added on day 3 and kept at a constant concentration (1 uM) for two days, a high frequency of the Tim3^−^TCF-1^+^ subset (48.5%) was maintained while the frequency of terminally exhausted (Tim3^+^TCF-1^−^) T cells was decreased to 17.1% (Day 3 is the initial stage of T cell exhaustion when ~95% T cells are still Tim3^−^TCF-1^+^, **Figure S2**). Furthermore, Taz treatment also decreased the expression of other inhibitory receptors (*e.g*., 2B4, CD39 and LAG3) but increased the expression of the memory marker CD62L (**Figure 2B**). Importantly, OT-I T cells treated with Taz exhibited comparable proliferation rate with cells cultured in normal T cell medium (**Figure 2C**). By day 7, Taz-treated OT-I T cells, upon restimulation, were less apoptotic than control, untreated T cells (**Figure S3A-C**). In addition, higher levels of multiple effector cytokines were detected in Taz-treated T cells compared to control T cells (**Figure S3D-G**). Consequently, enhanced killing of target 4MOSC1-SIINFEKL cells was observed (**Figure S3H**). Together, these observations indicate that Taz treatment maintains a progenitor-like phenotype in the cultured T cells by inhibiting their terminal differentiation without negative impact on T cell proliferation.

**Figure 2.**
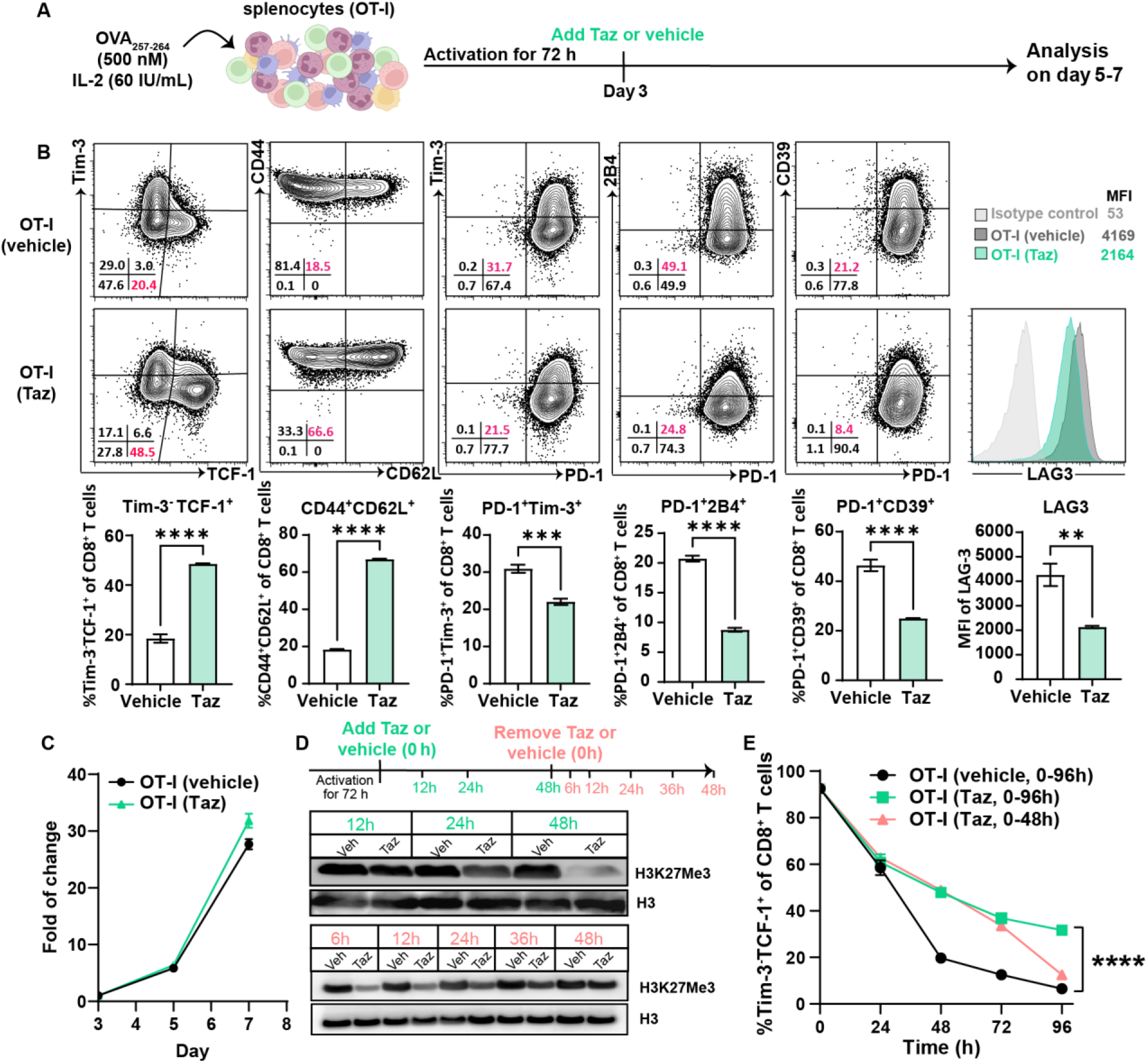
Transient inhibition of EZH2 by tazemetostat maintain progenitor-like features of OT-I cells. Phenotype analysis of OT-I T cells activated by OVA_257-264_ (500 nM) with 60 IU/mL IL-2 for 3 days and expanded in the presence or absence of Taz (1.0 uM) for 2 days (day3-5) (**A-B**). Cell proliferation of OT-I cells expanded w/ or w/o Taz (1 uM, Day3-7) (**C**). Western blot of histone proteins from OT-I T cells activated by OVA_257-264_ (500 nM) with 60 IU/mL IL-2 for 3 days and expanded in the presence or absence of Taz (1.0 uM) (**D**) for 12, 24, 48 h (upper panel), and then Taz was removed at 48h, OT-I T cells were collected after the removal of Taz at 6, 12, 24, 36, 48h (lower panel). Phenotype versus time analysis of OT-I T cells expanded in the presence or absence of Taz (**E**). Histone 3 (H3) was used as a reference protein. 3 repeats for each group (n=3), ^ns^P > 0.05; *P < 0.05; **P < 0.01; ***P < 0.001; ****P < 0.0001; analyzed by student T-test.

To track the kinetics of phenotypic and epigenetic changes of OT-I T cells expanded with Taz, cells were collected at different time points during the expansion course for Western blot and flow cytometry-based phenotype analysis. At approximately 24 h upon the addition of Taz, notable downregulation of H3K27Me3 was detected. By 48 h, the formation of H3K27Me3 was almost completely suppressed (**Figure 2D**), which was accompanied by a concomitant increase of the frequency of Tim3^−^TCF-1^+^ cells (**Figure 2E**). We observed that the Taz-induced inhibition of the EZH2 enzymatic activity was reversible such that at 36 h after the removal of Taz, H3K27Me3 was gradually recovered and reached a comparable level as cells expanded in a standard T cell medium in the absence of Taz (**Figure 2D**). Not surprisingly, upon the Taz removal, the frequency of Tim3^−^TCF-1^+^ OT-I T cells decreased to a similar level as their counterparts cultured without Taz (**Figure 2E**).

### Transient EZH2 inhibition improves *in vivo* proliferation, homeostasis, and recall response of transferred T cells

Based on the differences in the Tim-3 and TCF-1 expressions, OT-I T cells cultured with Taz would be endowed with progenitor-like properties, the hallmark of which is the enhanced capabilities of homeostatic proliferation and recall response. To evaluate these properties, OT-I splenocytes were first subjected to antigen stimulation, followed by the expansion with Taz or vehicle for 4 days. OT-I T cells (CD45.2+/+) treated with or without Taz were transferred into congenic C57BL/6J mice (CD45.1+/+ or CD45.1+/−) (**Figure S4A**) separately. At 24 h after adoptive transfer, the recipient mice were infected with *Listeria monocytogenes* (LM)-OVA (1×10^4^ CFU/mouse). Compared with OT-I T cells expanded in the absence of Taz, OT-I T cells expanded with Taz proliferated significantly better upon LM-OVA infection with twofold higher frequency detected in total CD8^+^ T cells in blood of the recipient mice on day 7 (**Figure S4B**) and 14 (**Figure S4C**), indicating a stronger recall response of Taz-treated OT-I CD8^+^ T cells. Higher frequencies of OT-I cells from the Taz-treated group were also found in the spleen (**Figure S4D**) and the liver (**Figure S4E**) on day14 after LM-OVA infection.

To determine the long-term survival of Taz-treated T cells, we transferred equivalent numbers of Taz-treated or vehicle-treated OT-I T cells (CD45.1+/−) to congenic C57BL/6J mice (CD45.2+/+) and harvested organs on day28 after the adoptive transfer to quantify the OT-I cell numbers. OT-I T cells pre-treated with Taz during the *in vitro* expansion showed improved homeostasis capabilities as two-threefold OT-I cells were found in spleen, liver, and lymph nodes (**Figure S5**). To further determine the recall capabilities, equivalent numbers (1×10^4^ cells/mouse) of Taz-treated or vehicle-treated OT-I T cells (CD45.2+/+) were transferred to congenic C57BL/6J mice (CD45.1+/−) (**Figure 3A**). The recipient mice were infected with LM-OVA (1×10^4^ CFU/mouse) on day 30 after adoptive transfer. After LM-OVA challenge, OT-I T cells pre-treated with Taz during *in vitro* expansion showed a stronger response evidenced by a higher ratio among total CD8^+^ T cells and more cell numbers were found in blood from day 7 to 14 (**Figure 3B, C**). On day30 after LM-OVA infection, more OT-I CD8^+^ T cells from Taz-pretreated group homed to the spleen and the liver as compared to the vehicle-treated group (**Figure 3D, E**). These data provided strong support that transient EZH2 inhibition via Taz-treatment during *ex vivo* expansion facilitates the production of T cells with better capabilities of homeostatic proliferation and recall response.

**Figure 3.**
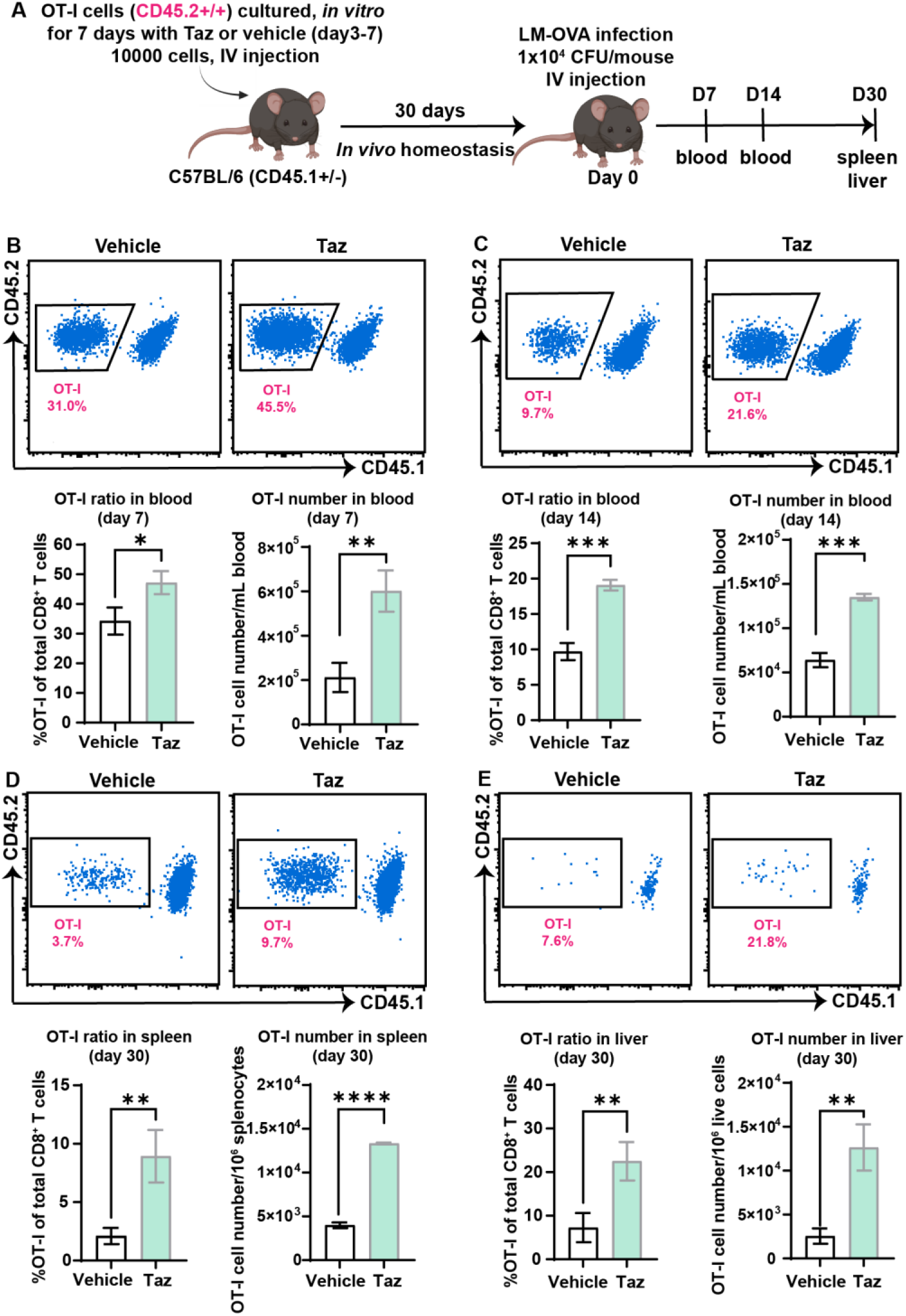
EZH2 inhibition by tazemetostat in OT-I cells during *in vitro* expansion improves *in vivo* homeostasis and response of OT-I cells for LM-OVA infection. OT-I splenocytes (CD45.2+/+) were activated with 500 nM OVA_257-264_ for 3 days and expanded in presence or absence of Taz (1 uM) for 4 days (day3-day7). 30 days after 1×10^4^ OT-I CD8^+^ T cells were intravenously transferred into C57BL/6 mice (CD45.1+/−), mice were infected with 1×10^4^ CFU LM-OVA (**A**). Ratio of OT-I cells in CD8^+^ T cells or exact number of OT-I cells in blood were analyzed on day 7 (**B**) and day 14 (**C**), in spleen (**D**) and liver (**E**) were analyzed on day 30. 3 repeats for each group (n=3), ^ns^P > 0.05; *P < 0.05; **P < 0.01; ***P < 0.001; ****P < 0.0001; analyzed by student T-test.

### Transient EZH2 inhibition synergies with checkpoint blockade to enhance anti-tumor efficacy of adoptively transferred T cells

We selected the aggressive, checkpoint blockade-resistant B16-OVA melanoma model to evaluate the activities of Taz-expanded T cells for treating solid tumors *in vivo*. OT-I T cells expanded with or without Taz were intravenously infused individually into C57BL/6 mice bearing established B16-OVA tumors, followed by anti-PD-1 administration on day14 and day 21(**Figure 4A**). In the control groups, mice were treated with PBS only (**Figure 4B, G**). Although OT-I T cells that were expanded without Taz alone or administered in a combination with anti-PD-1 only afforded modest tumor control (**Figure 4C-D, G**), OT-I T cells expanded with Taz significantly slowed down tumor growth with median tumor size being only 1/8 of that treated with vehicle-treated T cells on day 20 after adoptive T cell transfer (**Figure 4E, G**). Even better tumor control was realized when OT-I T cells that were expanded with Taz were used in a combination of anti-PD-1 (**Figure 4F-G**). Remarkably, whereas no mice in the control groups that received adoptive transfer of vehicle-treated OT-1 T cells survived up to day 24, >80% mice treated with Taz-expanded T cells were still alive by day 30 (**Figure 4H**). The median survival was extended for more than two weeks (from 22 to 37 days) for the mice treated with OT-I (w/ Taz) T cells plus PD-1 blockade as compared with that of mice received OT-I (w/o Taz) plus anti-PD-1 (**Figure 4H**).

**Figure 4.**
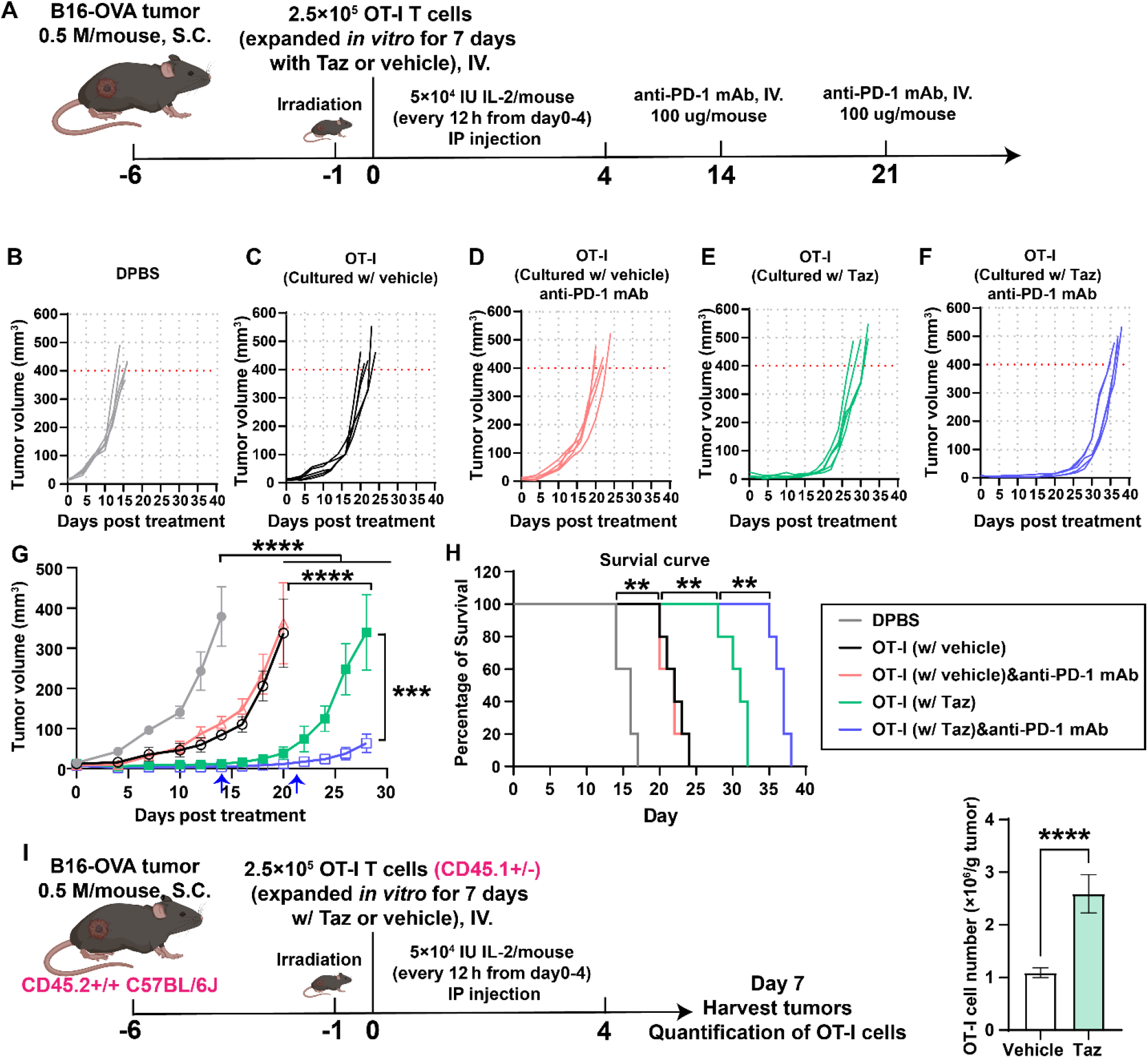
EZH2 inhibition by tazemetostat in OT-I cells during *in vitro* expansion improves *in vivo* efficacy and immune checkpoint response of OT-I cells against B16-OVA tumors. C57BL/6 mice were subcutaneously inoculated with B16-OVA tumor cells (6×10^5^). After 6 days (Day 0), OT-I T cells (2×10^5^/mouse), *in vitro* expanded with and without Taz (1 uM) for 4 days (day3-day7), respectively, were transferred, followed by IL-2 administration and anti-PD-1 antibody treatment on day 14 and day21 (**A**). In control groups, mice were administered with PBS buffer only. Tumor sizes (**B-G**, anti-PD-1 injections were indicated by blue arrows) and survival (**H**) were recorded every 2/3 days. OT-I cell numbers in tumors were analyzed on day 7 after adoptive cell transfer (**I**). 5 mice were included in each group (n = 5), ^ns^P > 0.05; *P < 0.05; **P < 0.01; ***P < 0.001; ****P < 0.0001, statistical analyses were performed by using one-way ANOVA followed by Tukey’s multiple comparisons test for tumor growth curves and log-rank test for survival curves.

To determine if better tumor control afforded by Taz-expanded OT-I T cells was caused by improved T cell tumor infiltration or survival, OT-I (CD45.1+/−) cells were intravenously injected into CD45.2+/+ mice bearing established B16-OVA tumors following exactly same procedures as described above (**Figure 4I**). On day 7 after the adoptive cell transfer, tumors were harvested for analysis. Approximately twofold higher numbers of OT-I T cells pretreated with Taz were found in tumors (**Figure 4I**). Taken together, these results indicated that compared to the vehicle-treated OT-I T cells, cells expanded with Taz possess markedly higher activities to control tumor growth *in vivo*. These Taz pre-treated cells consist of a higher fraction of less-differentiated, progenitor-like CD8^+^ T cells whose effector functions are successfully unleashed upon PD-1 blockade.

### Tazemetostat causes a gradual derepression of gene expression to induce a unique transcriptomic profile

To understand the molecular mechanism by which tazemetostat (taz) treatment redirects T cell fate away from terminal exhaustion and enhances the T cell function for adoptive cell therapy, we determined the transcriptomic profile of Taz or vehicle treated OT-I cells at 24 h and 48 h post treatment as described in Figure 1. Bulk transcriptomes of Taz-treated OT-I cells were similar to vehicle-treated controls at 24h post-treatment but diverged significantly at 48 h (**Figure 5A, B**). A higher number of genes was significantly upregulated than downregulated in the Taz group, as expected given EZH2’s role in silencing of gene expression (**Figure 5B**). Whereas 155 and 1847 genes were significantly upregulated in Taz treated cells at 24 and 48 h, respectively (adjusted *p*<0.05, fold change>2), 10 and 1213 genes were significantly downregulated at these timepoints (**Figure 5B**). The majority of differentially expressed genes (DEGs) at 24 h were also differentially expressed at 48 h (**Figure 5B**). These data suggest Taz causes major gene expression changes between 24-48 h post treatment. Examining DEGs by hierarchical clustering revealed groups of functionally related genes: Taz downregulated genes associated with MYC signaling and T cell stimulation such as *Srm* or *Nolc1* (**Figure 5C**, top cluster). *Myc* itself and some genes associated with effector differentiation such as *Gzmb* and *Ldha* were also downregulated (**Figure 5C**). By contrast, memory and naïve associated genes such as *Tcf7*, *Ccr7* and *Foxo1* were upregulated in the Taz group (**Figure 5C**). Among known EZH2 target genes, *Cdkn2a encoding two cell cycle regulators* was upregulated as expected (**Figure 5C**), and most EZH2-silenced memory-associated transcription factors^[23]^ were also upregulated (**Figure S6A**). One exception was *Eomes*, an EZH2-repressed gene downregulated after 48h of Taz treatment (**Figure S6A**).

**Figure 5.**
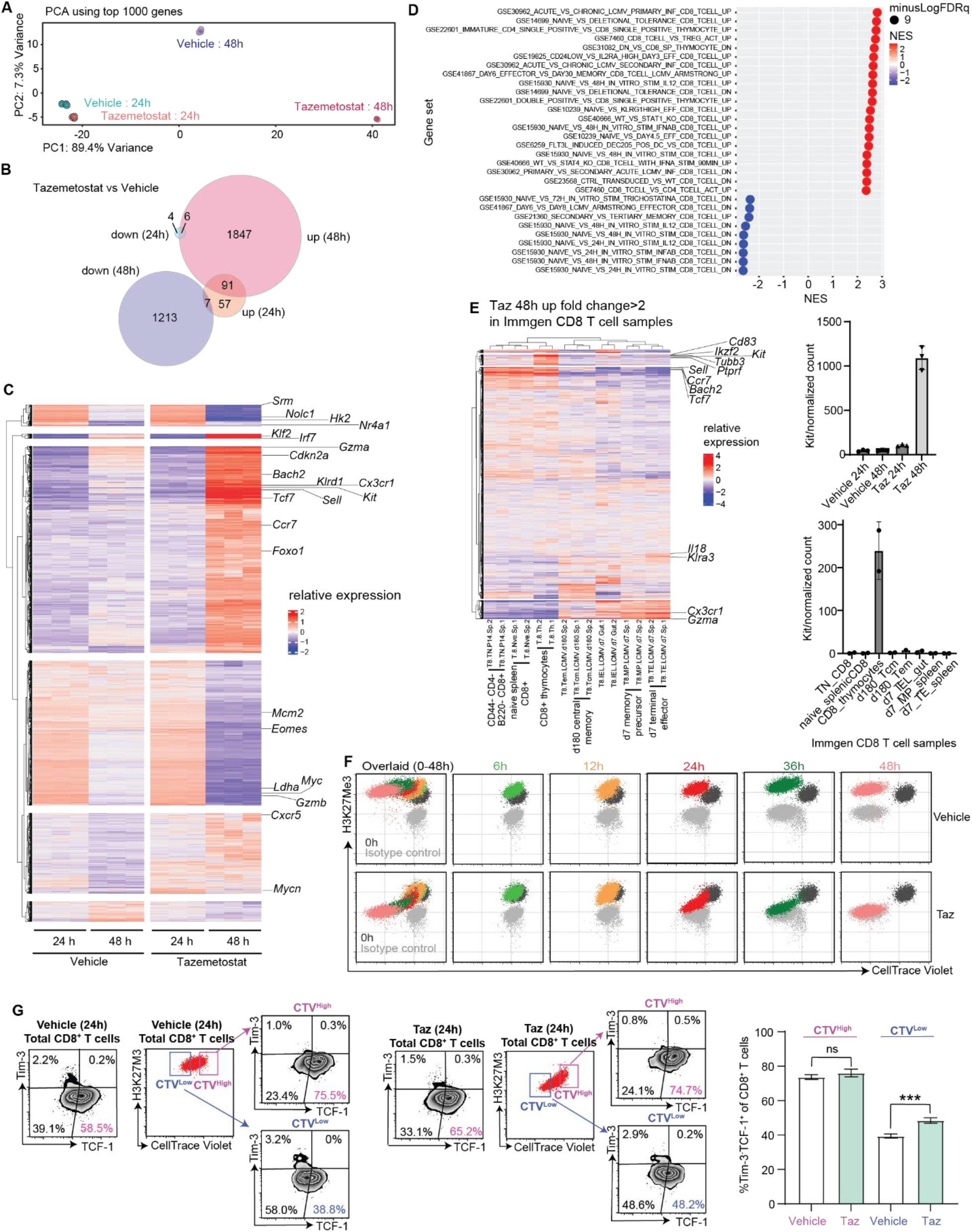
Transcriptomic profile of OT-I cells treated with Taz or vehicle. OT-I splenocytes were activated with treated for 24 or 48 h in the presence of Taz and subjected to RNA-seq. Principle component analysis (**A**); summary of differentially expressed genes, cut-off of adjusted *p*<0.05 was used (**B**); heatmap of the relative expression of all significant differentially expressed genes (**C**); gene set enrichment analysis of selected immune related gene sets at 48 h of taz vs vehicle treatment; (**D**) relative expression of taz-upregulated genes (fold change >2) in publicly CD8 T cell samples from Immgen (GSE109125) (**E**); OT-I cells were activated with 500 nM OVA_257-264_ for 3 days, labeled with CTV and cultured in the presence of Taz or vehicle for the durations indicated, flow cytometry analysis shown (**F**); expression of TCF1 and Tim3 in Taz or vehicle treated CTV^high^ and CTV^low^ T cells at 24 h post treatment (**G**). Statistical analysis of differential expression was performed by DESeq2, gene set enrichment analysis was performed by GSEA, Student’s two-sided t-test was used to compare samples with two groups. CTV, Cell Trace Violet; DEG, differentially expressed gene.

We performed gene set enrichment analysis to examine the impact of Taz-treatment on CD8^+^ T cells that are undergoing the initial stage of exhaustion using annotated gene signatures. Taz-treatment strongly suppressed MYC signaling, and mitosis associated signatures (**Figure S6B**, NES<-2) while enhancing TNF signaling via NF-kB (**Figure S6B**). Among transcription factor motifs, E2F motifs dominated those downregulated by Taz whereas FOXO1 motifs were enriched among upregulated genes (**Figure S6C**). In accordance with published studies that analyze the impact of EZH2 on CD8^+^ T cells during the early stage of acute LCMV infection,^[22, 23]^ gene sets enriched in Taz-treated T cells were upregulated in naïve T cells in comparison to effector and *in vitro* stimulated T cells (**Figure 5D**). Interestingly, despite the upregulation of memory/naïve genes and suppression of stimulation-associated gene sets, Taz also enhanced the expression of gene sets uniquely found in thymocytes (**Figure 5D**). To our surprise, gene sets upregulated in CD8 T cells from acute versus chronic LCMV infection (GSE30962) and day 6 effector vs day 30 memory CD8 T cells during acute LCMV infection (GSE41867) were also enhanced by Taz treatment. We then analyzed CD8 T cells RNA-seq data from the Immgen consortium in which various immune cell subsets were profiled using a highly standardized protocol. The expression pattern of Taz 48 h upregulated genes showed a strong cluster of naïve associated genes (**Figure 5E**). However, a gene cluster consisting of *Kit, CD83, and Ikzf2*, that is preferentially expressed in thymocytes and another cluster enriched in d7 effector CD8 T cells, exemplified by *Cx3cr1* and *Gzma*, were also observed (**Figure 5E**). Moreover, we also noticed that genes encoding pathogen-specific effector T cell secreted chemokines, including CCL3, CCL4, CCL5 and XCL1, were upregulated upon Taz treatment, which was further confirmed by antibody staining (**Figure S7**). Together, these observations demonstrate that Taz treatment does not simply increase the expression of memory/naïve associated genes, but rather generates a unique transcriptomic program with contributions from memory, naïve, effector and thymocyte associated genes.

Given that H3K27me3 histone levels change in a DNA replication-dependent manner and that H3K27 trimethylation level thresholds control gene expression (PMID: 30146316, 32027840), we hypothesized that Taz treatment might preferentially affect rapidly dividing T cells. To test this, OT-I T cells were stained with CellTrace Violet (CTV) after activation and cultured with Taz for 48h. As shown in **Figure 5F**, H3K27Me3 level gradually decreased during Taz treatment. At 24h of the treatment, CTV-high cells had comparable levels of H3K27Me3 with the untreated/pre-treatment cells whearas CTV-low cells approached the detection limit of H3K27Me3 by flow cytometry (corresponding to a nonspecific isotype control) (**Figure 5F**). Moreover, only CTV-low cells from Taz treatment group showed a higher proportion of Tim-3^−^TCF-1^+^ compared with untreated cells (**Figure 5G**). These data show that Taz differentially affects T cells depending on their rate of division and suggest that different phenotypic changes could be achieved depending on the duration of treatment.

### Preferential demethylation of promoter-associated H3K27 by tazematostat

Tazematostat is known to inhibit the ability of EZH2 to catalyze H3K27 methylation. Therefore, we examined the effect of *in vitro* Taz treatment on the landscape of H3K27me3 in OT-I T cells and asked whether the observed RNA expression changes could be attributed to changes in H3K27me3. Chromatin analysis using ChIP-seq identified many H3K27me3 peaks and the majority of them were affected by Taz treatment (**Figure 6A**). As expected, the large majority of Taz-affected H3K27me3 peaks had decreased abundance in Taz-treated cells (**Figure 6A**). There was a significant enrichment of peaks within promoter regions, especially within 1 kb of the promoter, within peaks decreased by Taz treatment (**Figure 6A**). Consistently, peaks within 1 kb of the transcription start site (TSS) were enriched in the Taz-decreased peak set compared to those unchanged with Taz (**Figure 6B**). The distribution of Taz-reduced TSS-associated peaks was skewed towards 3’ TSS-flanking peaks compared to the 5’ TSS-flanking sequence and Taz-reduced peaks were also enriched in exon-associated peaks (**Figure 6C, A**). As suggested in previous studies, we also observed a clear reduction in H3K27me3 associated with the start exon and promoter of *Tcf7* (**Figure 6D**).

**Figure 6.**
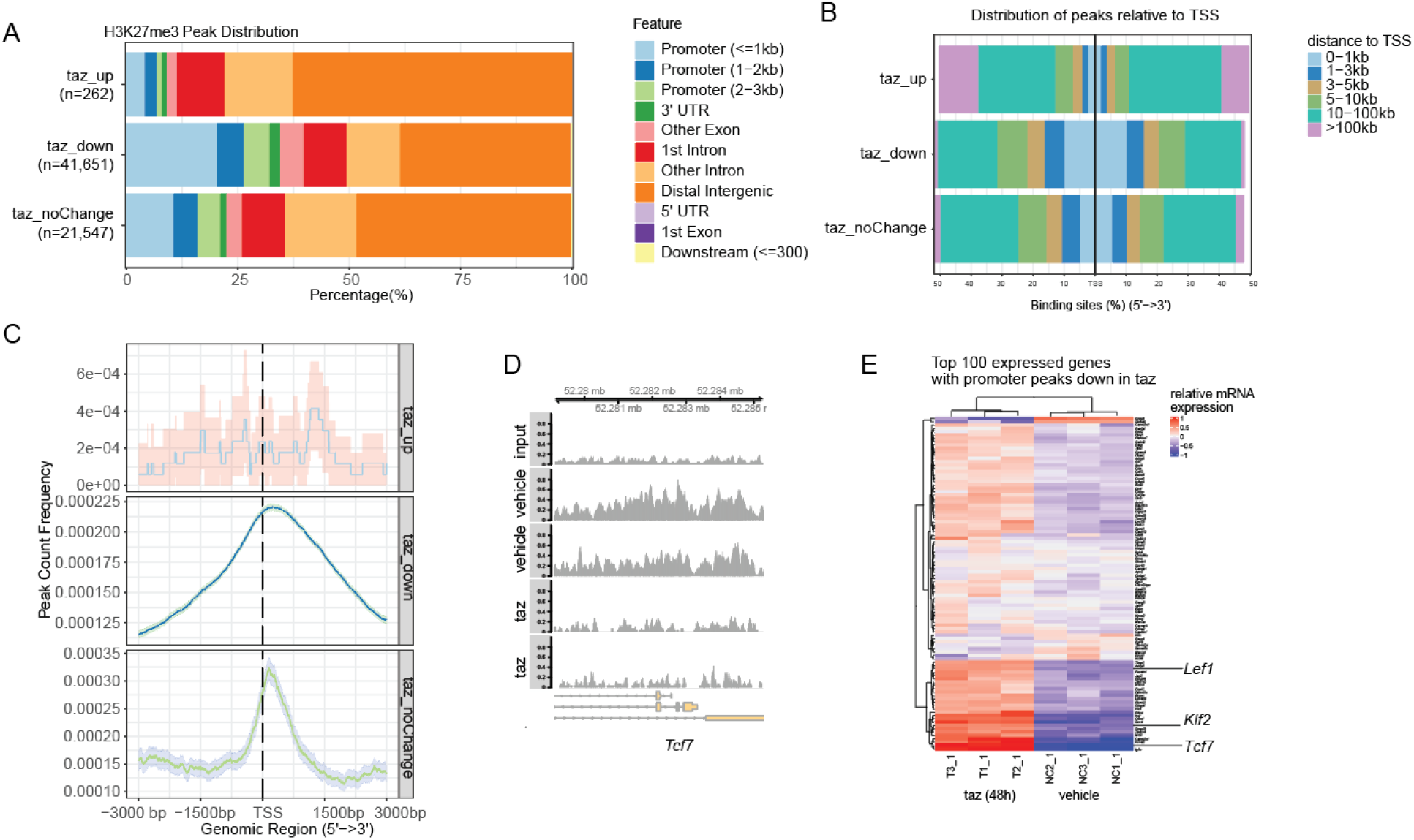
ChIP-seq identifies demethylation of promoter-associated H3K27 in Taz-upregulated genes. (**A**) Composition of H3K27me3 peaks by dominant genomic annotation, peaks grouped by the effect of Taz on their abundance; (**B**) composition of peaks by distance from transcription start site (TSS); (**C**) distribution of peaks within 6 kb of TSS grouped by Taz effect; (**D**) normalized distribution of peaks in the *Tcf7* promoter region; (**E**) relative mRNA expression of the top 100 protein-coding genes whose promoter-associated peaks were reduced by Taz. UTR, untranslated region; Taz, tazemetostat.

The majority of transcripts of genes whose promoter-associated peaks were reduced by Taz were upregulated compared to vehicle treated cells (**Figure S8**). Among the top Taz-reduced peaks were promoter peaks in the transcription factors *Tcf7*, *Klf2* and *Lef1*, confirming observations in T cells in other experimental models including LCMV.^[33]^ These results suggest that the increased expression of TCF1 at the protein and mRNA levels is accomplished by Taz by inhibiting the histone methylation of the promoter region.

### Transient inhibition of EZH2 reduces exhaustion features of antigen specific CD8+ T cells during LCMV Cl13 infection

T cell exhaustion caused by persistent antigen stimulation was first described in chronic virus infection in mice, and high dose *lymphocytic choriomeningitis* virus Clone 13 (LCMV Cl13) infection represents well-characterized murine models for chronic viral infections.^[34, 35]^ To substantiate the ability of Taz to reduce exhaustion and enhance stem-like features of exhausted CD8^+^ T cells, splenocytes from LCMV Cl13 infected mice were harvested at day15 post LCMV Cl13 infection and stimulated with T cell cognate peptides in the presence of Taz, vehicle or anti-PD-L1. Anti-PD-L1 antibody was used as a control in this assay, which increases the expression of inhibitory receptors during this stimulation.^[36]^ On day 5 post treatment, antigen-experienced T cells (CD44^+^) cultured with Taz showed significantly lower levels of inhibitory receptors expression, including TIM3 and LAG3, than vehicle-treated cells (**Figure 7A, B**). To determine the progenitor-like T cells, Ly108 was used as an alternative marker of TCF-1, which has been shown to be co-expressed with TCF-1.^[37]^ The percentage of progenitor-like Ly108^+^ Tim3^−^ cells of exhausted CD8 T cells was significantly higher in the Taz-treated group (**Figure 7C**), showing that Taz treatment induces antigen specific CD8^+^ T cells to be more progenitor phenotype. As with the OT-I experiment, the total number of CD8^+^ T cells, indicating the proliferation of antigen specific CD8^+^ T cells, was the same in Taz and vehicle treated wells (**Figure 7D**). Therefore, Taz can reduce markers of exhaustion and promotes progenitor-like cells in multiple mouse models without sacrificing the proliferation of antigen specific CD8^+^ T cells.

**Figure 7.**
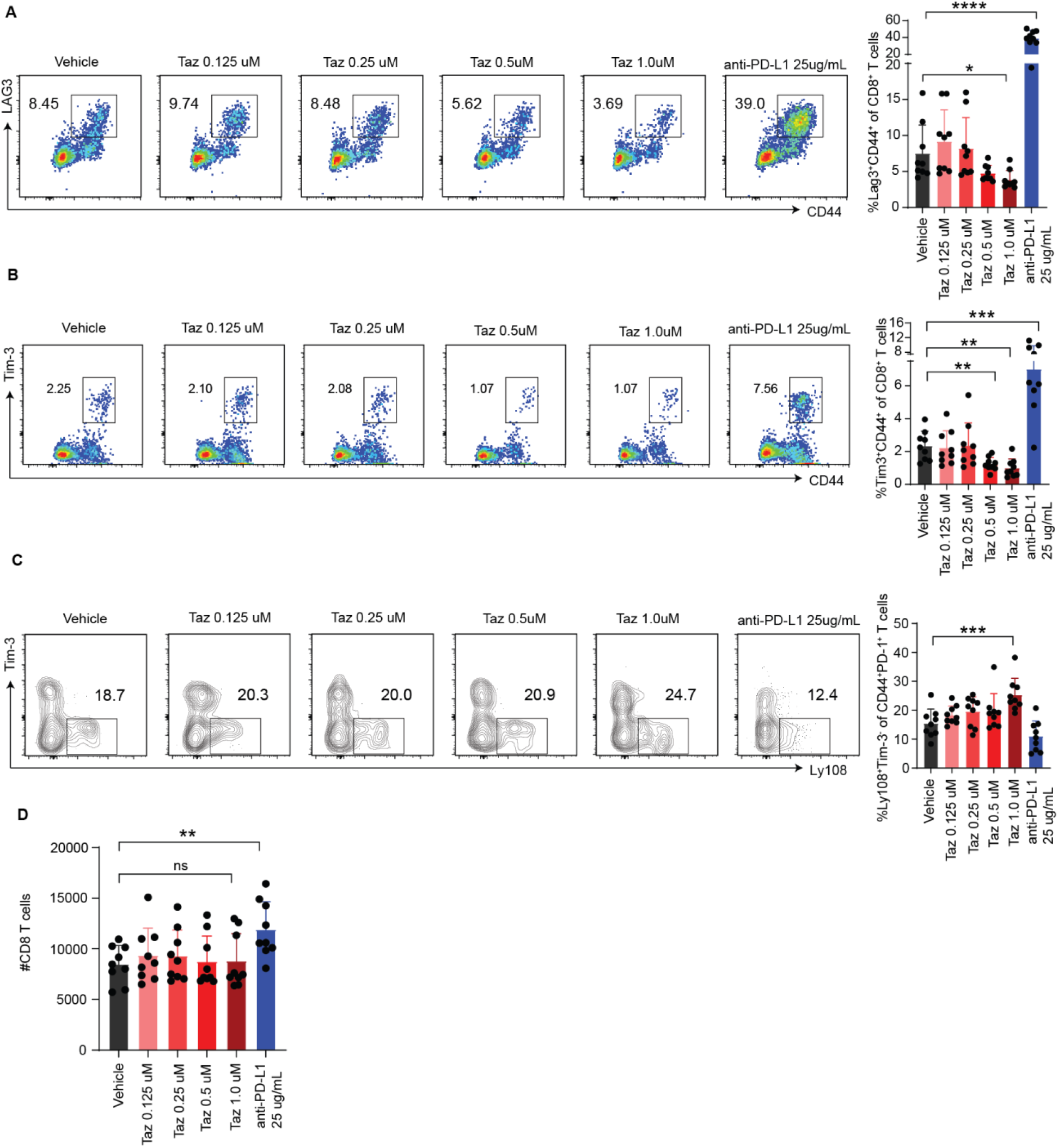
Transient inhibition of EZH2 with tazemetostat reduces exhaustion features of antigen specific CD8^+^ T cells. Splenocytes from C57BL/6J mice infected with LCMV Cl13 were harvested at d15 post infection and stimulated with viral peptides in the presence of Taz, anti-PD-L1 or vehicle. T cells were examined by flow cytometry at d5 of culture: analysis of LAG3 (**A**) and Tim-3 (**B**) expression on antigen-experienced T cells; (**C**) analysis of stem-like Ly108^+^ Tim-3^−^ T cells, gated on antigen-experienced exhausted T cells. (**D**) CD8^+^ T cell numbers. Statistical analysis was performed using one-way ANOVA and Dunnett’s post-test, ^ns^P > 0.05; *P < 0.05; **P < 0.01; ***P < 0.001; ****P < 0.0001.

### Transient EZH2 inhibition generates less-differentiated T cells in human PBMCs

After confirming that transient EZH2 inhibition by Taz could enhance the *in vivo* persistence and anti-tumor activities of murine CD8^+^ T cells upon adoptive transfer, we next sought to determine if *ex vivo* Taz treatment may alleviate T cell dysfunction and assist the generation of progenitor-like human T cells. A key factor to be evaluated was whether Taz-mediated EZH2 inhibition would undermine T cell activation and expansion. We adopted a modified chronic TCR stimulation assay,^[38]^ in which unfractionated human peripheral blood mononuclear cells (hPBMCs) were first stimulated with a mixture of irradiated hPBMCs from five HLA-mismatched healthy donors for 6 days followed by further stimulation with plate-bound anti-CD3. On Day 13, a majority of chronically stimulated CD8^+^ T cells entered a dysfunctional stage with the expression of multiple inhibitory receptors, including PD-1, Tim-3 and LAG3 (**Figure S9A, B**).^[39, 40]^ To examine whether temporary EZH2 inhibition could modulate the commitment of T cells to dysfunction, Taz was added to the co-culture at the beginning of stimulation and its concentration was maintained at 1 uM till day 13 (**Figure S9A**). At this stage, although CD8^+^ T cells were found to be less differentiated (**Figure S9B, C**), the cell proliferation was significantly reduced for all three donors (**Figure S9D**) as EZH2 is known to be required for T cell activation^[41]^. We then switched the addition of Taz (1 uM) on day3 after the T activation was finished and kept it at this concentration during the entire course of expansion (**Figure 8A**). On Day 13, compared with hPBMCs expanded in the absence of Taz, cells cultured with Taz harbor a two to threefold higher frequency and absolute numbers of Tim-3^−^TCF-1^+^ CD8^+^ T cells (**Figure 8B, E**) without notable differences in cell proliferation (**Figure 8D, G**). Moreover, expansion in the presence of Taz generated a higher frequency of CD45RA^+^CCR7^+^ (SCM: stem cell memory), CD45RA^−^CCR7^+^ (CM: central memory), and CD45RA^−^CCR7^−^ (EM: effector memory) CD8^+^ T cells, but lower ratio of CD45RA^+^CCR7^−^ (EMRA: effector memory cells re-expressing CD45RA) CD8^+^ T cells in the final cell product (**Figure 8C, F**, differentiation level: SCM<CM<EM<EMRA). These Taz-treated cells were also found to be poly-functional with a higher fraction capable of co-production of IFN-γ and IL-2 upon stimulation (**Figure 8H-K**). Together, these results demonstrate that including Taz in the human T cells expansion culture after T cell activation is capable of decoupling T cell proliferation from differentiation, alleviating exhaustion, and producing T cells with less differentiated phenotypes.

**Figure 8.**
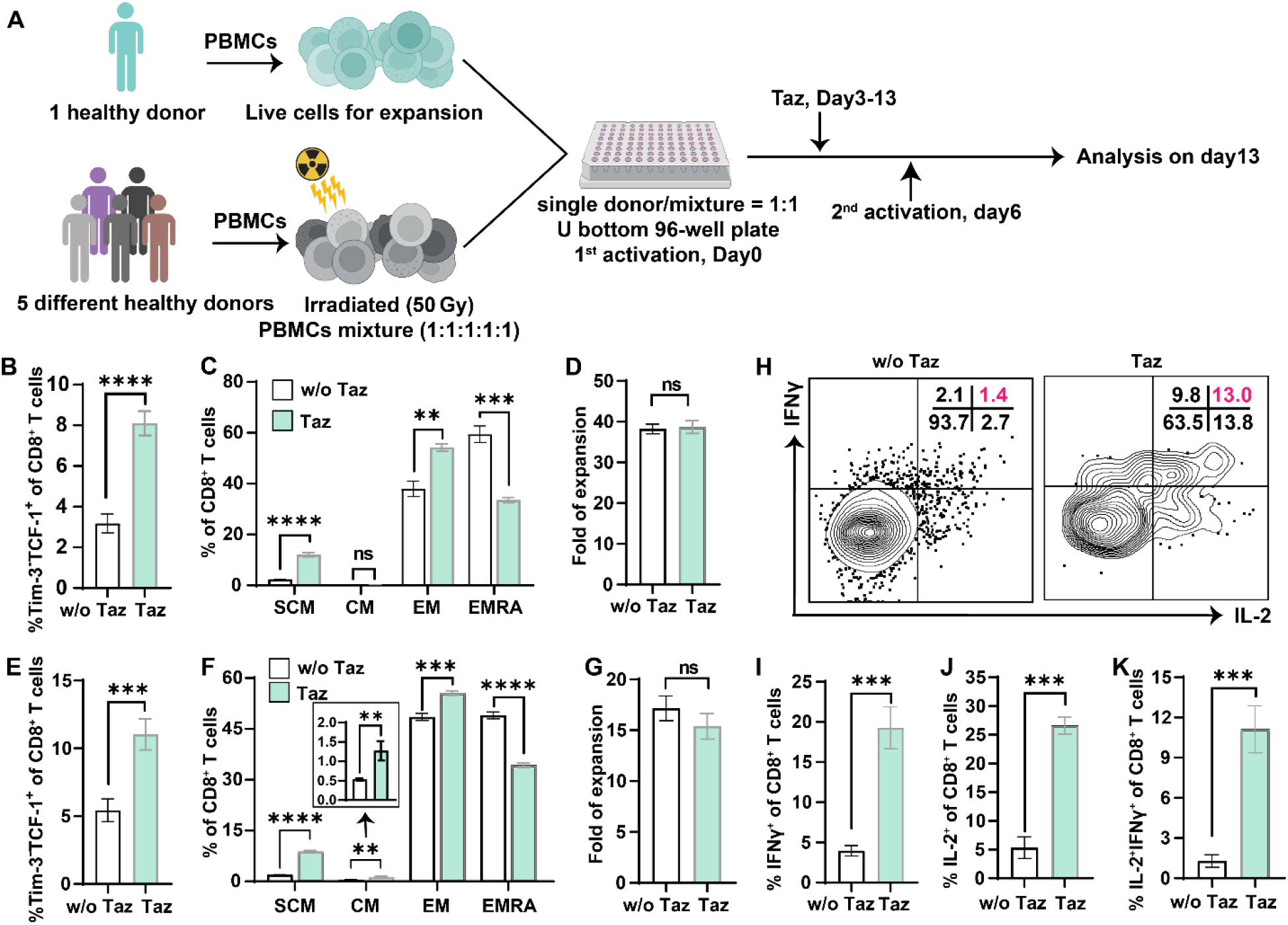
Tazemetostat enhances desired properties in human PBMCs. Human PBMCs were activated with an irradiated (50 Gy) mixture hPBMCs from another five different donors on day0 and day6 and expanded with 300 IU/mL IL-2 in the presence or absence of Taz (1uM, day3-day14) (**A**). Phenotype (**B-C** for donor A, **E-F** for donor B) and proliferation (**D** for donor A, **G** for donor B) analysis of CD8^+^ T cells on day 13 for 2 donors. SCM: CD45RA^+^CCR7^+^, CM: CD45RA^−^CCR7^+^, EM: CD45RA^−^CCR7^−^, EMRA: CD45RA^+^CCR7^−^, differentiation level: SCM<CM<EM<EMRA. Cytokine production of hPBMCs were cultured w/ or w/o Taz (1 uM, day3-day13) and activated with coated anti-CD3 antibody (1 ug/mL) and suspended anti-CD28 antibody (1 ug/mL) for 5 h (**H-K**). 3 repeats for each group (n=3), ^ns^P > 0.05; *P < 0.05; **P < 0.01; ***P < 0.001; ****P < 0.0001; analyzed by student T-test.

### Transient EZH2 inhibition alleviates CAR-T cell exhaustion

With the validation that Taz treatment could decouple T cell differentiation from proliferation when added following human T cell activation, we investigated if Taz could be used to alleviate exhaustion of genetically engineered CAR-T cells during *ex vivo* expansion. Exhaustion is a major obstacle that limits the efficacy of CAR-T therapy in both liquid and solid malignancies. CAR-T cells, e.g., CD22-m971-28z, ErbB2-28z, GD2-28z and HA-28z CAR-T cells, are known to develop varying degrees of exhaustion features during *ex vivo* expansion. T cells engineered to express the high-affinity anti-disialoganlioside (GD2) CAR HA-28z ^[42]^ develop functional, transcriptomic, and epigenetic hallmarks of exhaustion by day 11 after activation as a result of antigen-independent receptor clustering and tonic signaling.^[43]^ To test the effect of Taz, we transduced PBMCs from a healthy donor with HA-28z and cultured the sorted CD8^+^ CAR-T cells in the presence of Taz till day 20 (**Figure 9A**). At this time point, compared to CAR-T cells cultured in normal medium, cells cultured with Taz exhibited decreased inhibitory receptor (PD-1, LAG3, Tim-3) expression (**Figure 9B, D**) and a concomitant increase of TCF-1 expression (**Figure 9B, C**). During the entire course of expansion, no obvious impact on cell proliferation was observed upon the addition of Taz (**Figure 9E**). Notably, we found that a greater fraction of the Taz-treated CAR-T cells had intrinsic IL-2 production (**Figure 9F**), which is an unique feature of stem-cell-like CD8^+^ T cells.^[44]^ Likewise, a 2-5fold greater proportion of these Taz-treated cells were more polyfunctional, as measured by IFN-γ, TNF-α and IL-2 triple or double positive cells. Finally, the Taz-treated HA-28z CAR-T cells were found to be more cytolytic toward the Nalm6 target cells that were engineered to overexpress GD2 (Nalm6-GD2).

**Figure 9.**
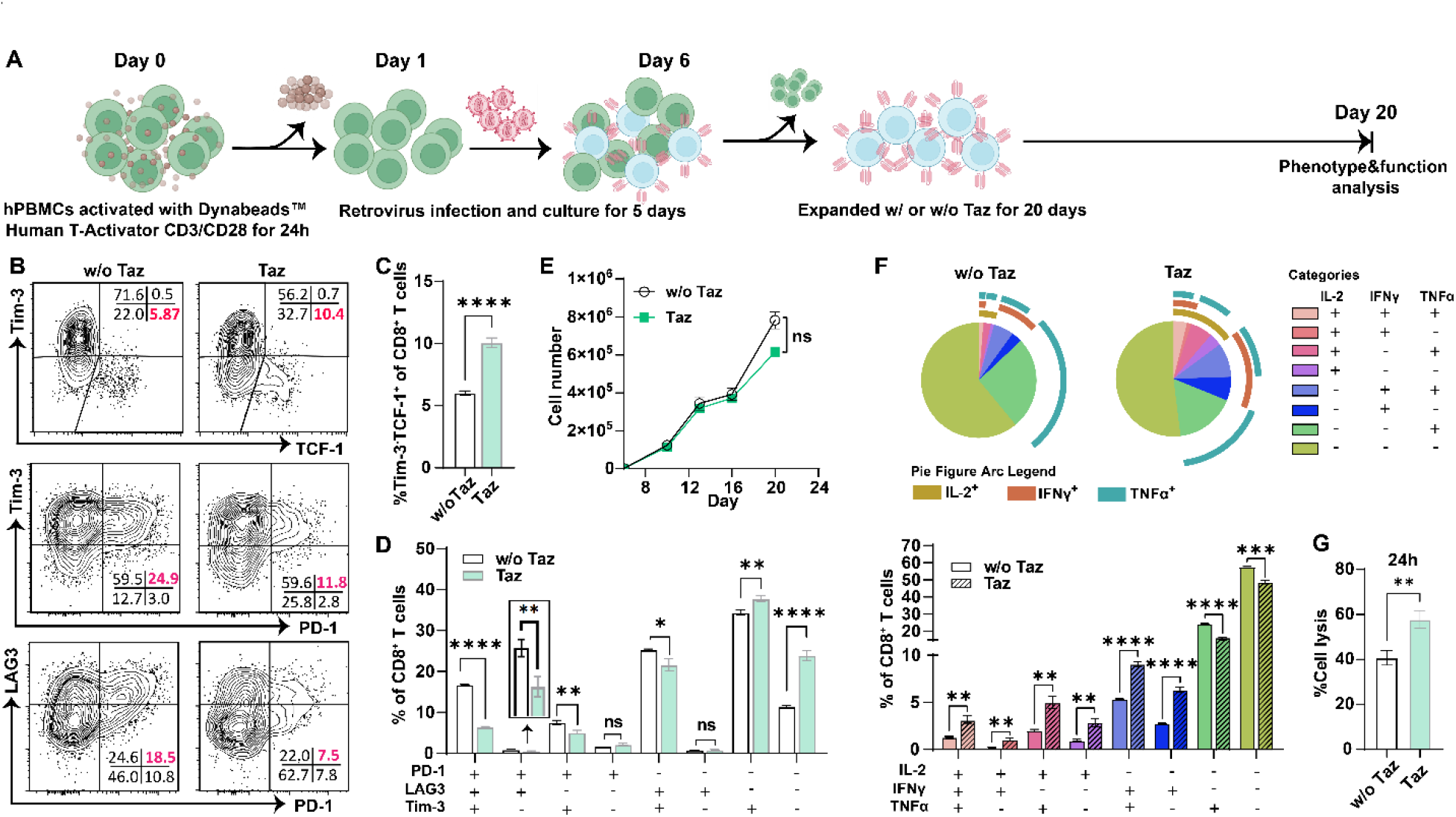
Tazemetostat decreases exhaustion phenotypes of HA-28z CAR T cells. Human PBMCs were transinfected with HA-28z and CD8^+^HA-28z CAR^+^ T cells were sorted for *in vitro* expansion with 150 IU/mL IL-2 in the presence or absence of Taz (1 uM) (**A**), phenotype analysis(**B-D**) and cell growth curve (**E**) on day 20 after activation. Cytokine production (**F**) and cell lysis (**G**) of HA-28z CAR-T cells stimulated or co-cultured with targeting cells Nalm6-GD2. 3 repeats for each group (n=3), ^ns^P > 0.05; *P < 0.05; **P < 0.01; ***P < 0.001; ****P < 0.0001; analyzed by student T-test.

## Discussion

Dynamic regulation of epigenetic and chromatin states influence how T cells acquire or lose plasticity and how particular T cell developmental trajectories are settled. As uncovered by Kaech and coworkers, EZH2-mediated epigenetic silencing is required for the differentiation of effector CD8^+^ T cells but not required for memory cell formation.^[22]^ Both *Ezh2*-deficient naïve and *in vivo* primed CD8^+^ T cells failed to make a strong *in vivo* response to viral infection or restimulation due to cell apoptosis and exhibited an impaired ability to produce inflammatory cytokines for effective pathogen clearance.^[41]^ Notably, if the deletion of *Ezh2* in CD8^+^ T cells occurred after activation, only cell cycle was delayed without increased apoptosis during acute viral infection. Consistent with this discovery, Zou et al. found that EZH2^+^CD8^+^ T cells were polyfunctional, apoptosis-resistant, and positively correlated with improved survival in ovarian cancer patients.^[45]^ Accordingly, *Ezh2* knockdown in human T cells elicited poor antitumor immunity. However, the functional role of EZH2 in T cell exhaustion is just started to be explored.

Recently, using a mouse LCMV infection model, Chang et al. discovered that in comparison to CD8^+^ T cells that had undergone their first division in response to LCMV-Arm, CD8^+^ T cells that had undergone their first division in response to LCMV-Cl13 expressed higher levels of EZH2.^[46]^ Examining this and additional public human and mouse bulk and single-cell RNA seq datasets, we observed that rather than being associated with a particular differentiation stage, the expression of EZH2 is tightly correlated with T cell proliferation —actively dividing cells of diverse phenotypes harbor the highest levels of *Ezh2* expression.

Because proliferative CD8^+^ T cells, including those giving rise to terminally exhausted cells, have high *Ezh2* levels and EZH2 is required for the effector function, we explored the feasibility of temporarily blocking its enzymatic activities by using tazemetostat (Taz), an inhibitor that is currently used for treating sarcoma.^[30]^ Pharmacological inhibition of EZH2 with Taz in T cells in the early stage toward exhaustion delayed their dysfunctional progression and facilitated the generation of progenitor-like T cells with polyfunctionality by reprogramming their epigenetic program. Critically, unlike most small-molecule modulators used previously to enhance T cell persistence and longevity,^[47]^ temporary treatment with Taz had no obvious impact on cell proliferation. Not only did Taz treatment assist in maintaining high TCF-1 expression, but it also facilitated T cells to produce a diverse array of chemokines and cytokines crucial for T cell proliferation, DC attraction, and tumor control. Rather than enhanced the expression of memory-associated genes, Taz induced a transcriptomic profile partially overlapping with those of thymocytes, naïve, memory and effector T cells. Interestingly, we found that Taz-mediated changes required T cell division, as nondividing T cells maintained normal H3K27Me3 and TCF-1 levels in the presence of Taz. This observation links the preferential expression of EZH2 in actively dividing cells to the selective impact of Taz on dividing T cells. Controlling how many divisions T cells undergo in the presence of Taz could therefore offer a strategy to avoid the detrimental effects of genetic *Ezh2* deletion.^[20, 22]^ Excitingly, adoptive transfer of CD8^+^ T cells expanded *ex vivo* with Taz followed by checkpoint blockade with anti-PD-1 afforded superior tumor control in a mouse model of melanoma.

Mackall and coworkers previously discovered that transient rest restores anti-tumor immunity in exhausted HA-28z CAR-T cells through epigenetic reprogramming. Treating Day 11-15 rested CAR-T cells did not compromise CAR-T cell viability, proliferation, and other functions, nor did it alter the rest-associated decrease of the expression of cell-surface inhibitory receptors.^[43]^ By contrast, in the current study, when HA-28Z CAR-T cells were treated with Taz starting on day 5 upon activation, alleviated exhaustion and augmented effector functions were observed from day12-20. These findings suggest that in the early stage along the pathway toward exhaustion there seems to be a convergence of epigenetic mechanisms that govern T cell phenotype and dysfunction.

As an oncogene, EZH2 promotes cell survival, proliferation, epithelial to mesenchymal, invasion, and drug resistance of cancer cells. Given its clear gain-of-function contributions to cancer and the capability of directly inhibiting its enzymatic function, EZH2 is an actively pursued target for anti-cancer therapy.^[48]^ On the other hand, EZH2 expression is also elevated in tumor-infiltrating regulatory T cells (Tregs). Bluestone and coworkers discovered that pharmacological inhibition of EZH2 using CPI-1205, an orally available selective inhibitor of EZH2, destabilized FOXP3 expression and impede tumor growth in murine tumor models.^[49]^ In an independent study led by Sharma, checkpoint blockade using the anti-cytotoxic T lymphocyte-associated protein 4 (CTLA-4) antibody ipilimumab was found to increase the EZH2 expression in human T cells across various tumor types.^[50]^ Increased EZH2 expression in T cells inversely correlated with clinical outcomes in a cohort of patients with prostate cancer. In murine tumor models, the combination use of CPI-1205 significantly improved antitumor responses elicited by anti–CTLA-4. These discoveries, combined with our findings, suggest the possibility of a potential clinical trial course in patients with primary or adaptive resistance to anti–CTLA-4 therapy, in which CPI-1205 plus ipilimumab would be administered first, followed by the administration of anti-PD-1 and the adoptive transfer of TCR-T or CAR-T cells that are expanded *ex vivo* with an EZH2 inhibitor.

## Acknowledgement

This work was supported by the NIH (R01AI143884 to J.R.T and P.W.). J. Z. is supported by the Cancer Research Institute/Irvington postdoctoral fellowship. We thank Prof. Crystal L. Mackall for the plasmid of HA-28z CAR and Nalm6-GD2 cells.

## Declaration of interests

The authors declare no competing interests.

## Methods

### Mice and cell line

C57BL/6J (both WT and CD45.1+/+ congenic strains) mice were purchased from Jackson Laboratory. OT-I mice (strain B6.*129S6*-*Rag2^tm1Fwa^*Tg(TcraTcrb)1100Mjb, CD45.2+/+) were purchased from Taconic Biosciences. OT-I+/− CD45.1+/− mice were bred in-house using C57BL/6J (CD45.1+/+) and OT-I mice (CD45.2+/+). All mice were bred and housed in specific pathogen free (SPF) rooms. All animal studies were approved by TSRI Animal Care and Use Committee. B16-OVA cells (provided by Prof. Gregoire Lauvau’s lab) and HEK293 Phoenix Ampho Packaging Cells (provided by Prof. Andrew Ward) were cultured in complete DMEM medium (Gibco, Cat#10569-010) supplemented with 10% fetal bovine serum (FBS, Hyclone, Cat#SH30088.03) and 1% penicillin-streptomycin (Gibco, Cat#15140-122). Nalm6-GD2 cells (provided by Prof. Crystal L Mackall’s lab) were cultured in RPMI1640 medium (Gibco, Cat#61870-127) supplemented with 10% FBS and 1% penicillin-streptomycin. 4MOSC1-SIINFEKL-GFP cells (provided by prof. J. Silvio Gutkind’s lab) were cultured in Defined Keratinocyte-SFM medium supplemented with EGF Recombinant Mouse Protein (5 ng/ml), Cholera Toxin (50 pM) and 1% antibiotic/antimycotic solution.

### Human blood samples

All human blood samples were purchased from the Scripps General Clinical Research Center (GCRC) with informed consent of anonymous healthy donors via Scripps Research’s Normal Blood Donor Service (NBDS) under a Scripps Institutional Review Board (IRB) protocol.

### *In vitro* activation of OT-I cells (both CD45.1+/− and DC45.2+/+)

For the comparation of OT-I cells activated with different concentrations of OVA_257-264_ (SIINFEKL, GenScript), OT-I splenocytes were activated with OVA_257-264_ (1 nM) for two days in T cell medium (RPMI1640 medium supplemented with 10% FBS, 1% penicillin-streptomycin, sodium pyruvate (1 mM, Gibco, Cat#11360-070), HEPES (10 mM, Gibco, Cat#15630-080), NEAA (1×, Gibco, Cat#11140 −050), 2-mercaptoethanol (Gibco, Cat#21985-023)) and expanded for another 5 days in T cell medium containing 60 IU/mL of IL-2 (Recombinant human IL-2, Teceleukin obtained from NIH), or OT-I splenocytes were activated with OVA_257-264_ (500 nM) for three days in T cell medium containing 60 IU/mL of IL-2 and expanded for another 4 days in the same medium. Cell numbers were recorded, and phenotype was analyzed at predetermined time points.

For the treatment of OT-I cells with Tazemetostat (Selleckchem), OT-I splenocytes were activated with OVA_257-264_ (500 nM) for three days in T cell medium containing 60 IU/mL of IL-2 and expanded in the same medium with or without Tazemetostat (1 uM) for another 2-4 days. Cells were harvested at predetermined time points for *in vitro* analysis and *in vivo* adoptive transfer.

### Western blot

Western blot analysis was performed using standard protocols. Histone proteins were isolated using a Histone Extraction Kit (Abcam, Cat#ab113476) following the manufacturer’s instructions and separated by 4-12% SDS-PAGE (Invitrogen, NuPage, Cat#NP0322BOX) in a 1×MES buffer (Invitrogen, Cat#NP0002). Transmembrane was performed using a Trans-Blot○R Turbo Transfer System (Bio-rad). The membranes were blocked for 1hr at room temperature with 5% non-fat milk (Research Products International, Cat#M17200-1000.0) in 1×Tris-buffered saline Tween-20 (TBST, G-Biosciences, Cat#R042), and analyzed with the indicated primary antibodies (biotinylated anti-H3K27Me3 (1:1000, Cell Signaling Technology, Cat#4395S, clone: C36B11), anti-H3 (1:1000, Cell Signaling Technology, Cat#4395S, clone: C36B11)) at 4 °C overnight. The membranes were washed with 1×TBST for three times followed by incubation with a streptavidin-horseradish peroxidase (1:2000, streptavidin-HRP) or an anti-rabbit antibody conjugated with HRP (1:2000, Sigma, Cat#12-348) at room temperature for 1 hr. The membranes were visualized using SuperSignal™ West Pico PLUS Chemiluminescent Substrate (Thermo Scientific, Cat#34580) by machine (Bio-rad) after washed with 1×TBST for three times.

### Flow cytometry analysis

For surface marker staining, single cell suspension in 1×DPBS (Gibco, Cat#14190-144) containing 2%FBS and 1 mM EDTA (FACS buffer) was collected into V-bottom plates, blocked with anti-mouse CD16/32 antibody (Biolegend, Clone: 93) for 10 min at room temperature, and incubated with indicated antibodies on ice for 30 min. Cells were washed with 1×DPBS for 2 times and stained with Ghost Dye™ Violet 510 (1:1000, Tonbobio) for 10 min in 1×DPBS at room temperature. Cells were then washed with FACS buffer and resuspended in the same buffer for flow cytometry analysis. For intracellular cytokine or intranuclear transcription factors staining, cells were first stained for surface markers and Ghost Dye™ Violet 510 as described above. Next, cells were fixed and permeabilized with a cytokine staining buffer set (Biolegend, Cat#420801 and Cat#421002) for cytokine/chemokine staining and a Foxp3/Transcription Factor Staining Buffer Set (eBioscience/Thermo Fisher Scientific, Cat#00-5523-00) for intranuclear staining following the manufacturer’s instructions. Human samples were also stained following similar procedures with antibodies that are specific to human markers.

### Antibodies for flow cytometry or CAR-T cell sorting

The following antibodies were purchased from Biolegend for mouse samples: CD16/32 (Clone: 93), CD8a (Clone: 53-6.7), PD-1 (Clone: 29F.1A12), LAG3 (Clone: C9B7W), Tim-3 (Clone: RMT3-23), 2B4 (Clone: m2B4 (B6)458.1), CD39 (Clone: Duha59), CD44 (Clone: IM7), CD62L (MEL-14), CD45.1 (Clone: A20), CD45.2 (Clone: 104).

The following antibodies were used for chemokine staining: CCL3 (MIP-1 alpha) Monoclonal Antibody (DNT3CC)-PE (eBioscience, Cat#12-7532-80), Mouse CCL4/MIP-1 beta Antibody (R&D, Cat#AF-451-SP), PE/Cyanine7 anti-mouse CCL5 (RANTES) Antibody (Biolegend, Cat#149105), Mouse XCL1/Lymphotactin Antibody (R&D, Cat#AF 486-SP), Donkey anti-Goat IgG (H+L) Cross-Adsorbed Secondary Antibody, Alexa Fluor™ 488 (Invitrogen, Cat# A-11055). CD8a-Pacific orange (Invitrogen, Clone: 5H10) antibody was purchased from Invitrogen/Thermo Fisher Scientific, and TCF-1 (AF488, Clone: C63D9) antibody was purchased from Cell Signaling Technology.

The following antibodies were purchased from Biolegend for human samples: Fc blocker (Cat#422302), CD8a (Clone: HIT8a), CD4 (Clone: A161A1), PD-1 (Clone: EH12.2H7), LAG3 (Clone: 7H2C65), Tim-3 (Clone: F38-2E2), CCR7 (Clone: G043H7), CD45RA (Clone: HI100), IL-2 (Clone: MQ1-17H12), IFNγ (Clone: 4S.B3), TNFα (Clone: MAb11).

Fluorescein (FITC) AffiniPure Sheep Anti-Mouse IgG, F(ab’)_2_ fragment specific (Jackson ImmunoResearch, Cat#515-095-072) was used for CAR-T cell sorting.

All samples were analyzed by CytKick™ Max Flow Cytometer.

### Intracellular cytokine production of OT-I cells

OT-I T cells were restimulated with anti-mouse CD3 antibody (Clone: 17A2, Biolegend, Cat#100238, 1 μg/mL, 100 uL/well, precoated onto the plate) and anti-mouse CD28 antibody (Clone: 37.51, Biolegend, Cat#102116, 1 μg/mL in medium) for 5-7h in the presence of Brefeldin A (Biolegend, Cat#420601) in a 96-well plate. Cell staining followed the protocol described above.

### OT-I killing assay against 4MOSC1-SIINFEKL cells

4MOSC1-SIINFEKL expressing GFP cells (10000 cells/well) were plated into a 96-well plate precoated with collogen, and co-cultured with OT-I cells at various effector/targeting cells ratio (2:1, 1:1, 0.5:1) for 20h. The fluorescence intensity of GFP was recorded by an Agilent BioTek Synergy H4 Hybrid Microplate Reader.

### *Listeria monocytogenes*-OVA (LM-OVA) infection

For the stimulation once OT-I cell was transferred, 5000 of OT-I cells (CD45.1+/−) *in vitro* expanded with or without Tazmezostat (1 uM) were transferred into gender matched congenic mice (CD45.2+/+, 7-week-old) via intravenous (IV) injection. 24h after the adoptive transfer, these mice were infected with LM-OVA (1×10^4^ colony-forming units (CFU)/mouse) by IV injection. On day7 and day14 post LM-OVA infection, blood was collected for analysis. Liver and lung were collected and homogenized for analysis on day14.

For the homeostasis study, OT-I cells (CD45.2+/+, 1×10^4^/mouse) were transferred into gender matched congenic mice (CD45.1+/−, 7-week-old) via IV injection. 30 days after the adoptive transfer, the mice were infected with LM-OVA (1×10^4^ CFU/mouse) by IV injection. OT-I cell numbers and frequencies in blood were analyzed on day 7 and day 14 after the infection. OT-I cell numbers and frequencies in liver and lung were analyzed on day 30 after the infection.

### Adoptive OT-I cell transfer for B16-OVA tumor treatment

6-7-week-old female C57BL/6J mice were subcutaneously implanted with 5×10^5^ B16-OVA cells (day −6). On day 5 (day −1) after the inoculation of B16-OVA cells, the mice were irradiated with a dose of 5 Gy for lymphodepletion. OT-I cells (2.5×10^5^ cells/mouse) from a female OT-I mouse *in vitro* expanded with or without Tazemetostat were transferred into the mice bearing B16-OVA tumors (day 0) followed by IL-2 (5×10^5^ IU suspended in 200 uL DPBS for each mouse) infusions every 12h for 4 days via intraperitoneal (IP) injection. Anti-PD-1 antibody infusions were performed on day 14 and day 21 after the adoptive transfer of OT-I cells with a dose of 100 ug/mouse by IV injection.

### B16-OVA tumor infiltrating OT-I cells analysis

6-7-week-old female C57BL/6J mice (CD45.2+/+) were subcutaneously implanted with 5×10^5^ B16-OVA cells (day −6). On day 5 (day −1) after the inoculation of B16-OVA cells, the mice were irradiated with a dose of 5 Gy for lymphodepletion. OT-I cells (2.5×10^5^ cells/mouse) from a female OT-I mouse (CD45.1+/−) *in vitro* expanded with or without Tazemetostat were transferred into the mice bearing B16-OVA tumors (day 0) followed by IL-2 (5×10^5^ IU suspended in 200 uL DPBS for each mouse) infusions every 12h for 4 days via intraperitoneal (IP) injection. Tumors were harvested on day 7 after the adoptive transfer of OT-I cells and homogenized to a single cell suspension for analysis.

### RNA-seq

OT-I splenocytes were activated with OVA_257-264_ (SIINFEKL, 500 nM) with 60 IU/mL of IL-2 for three days to obtain CD8^+^ T cells. The cells were further cultured with or without 1 uM Taz for 24 or 48 h before they are harvested for RNA extraction with the Arcturus™ PicoPure™ RNA Isolation Kit (KIT0204, ThermoFisher Scientific). Traces of genomic DNA were removed using RNase-Free DNase Set (79254, QIAGEN) following the on-column digesting protocol of the RNA isolation kit. Library construction and RNAseq data acquisition was done by the Novogene Corporation (Sacramento, CA) using Illumina HiSeq platform and PE150 strategy. Quality control of raw reads was performed using fastQC v0.11.9 (http://www.bioinformatics.babraham.ac.uk/projects/fastqc). Reads were aligned to the GRCm38-mm10 genome and gene counts calculated using STAR v2.7.6a. Normalization and differential expression analyses were performed using DESeq2 v1.38.1^[51]^. Heatmaps were plotted using the ComplexHeatmap package^[52]^. Gene set enrichment analysis was performed using GSEA^[53]^. Data from ImmGen dataset GSE109125 were used for the analysis of gene expression in mouse T cells. For the analysis of The Cancer Genome Atlas data, normalized RNA-seq values were used and transcriptome-wide Spearman rank correlation coefficients calculated for each tumor type using cBio. R v4.2.2 was used for the analysis and plotting of bulk RNA-seq data.

### Public Single-cell RNA-seq and microarray analysis

The public scRNA-seq datasets were analyzed with the Single Cell Portal (https://singlecell.broadinstitute.org/single_cell). Datasets from the following studies were used: PMID: 35549406, PMID: 35869073, PMID: 34493872, PMID: 32790115, PMID: 35995567, https://doi.org/10.1101/2020.03.31.018739. Public microarray data from CD8 T cells in LCMV infection from GSE30431 were analyzed using GEO2R.

### ChIP-seq

OT-I splenocytes were activated with 500 nM OVA_257-264_ for three days and then expanded in the presence of Taz (1 μM) or vehicle with IL-2 (60 IU/mL) for 36h. Dead cells were removed by ficoll and live cells were collected and snap-freezed. Samples were sent to Active Motif for DNA preparation, chromatin Immunoprecipitation and sequencing. Raw reads were aligned to the mm10 genome using STAR v2.7.6a. Peak counting and differential abundance analysis were performed using csaw v1.32.0.^[54]^ The input control was used to establish background peak abundance. Peaks with falls discovery rate < 0.05 were considered significant in the differential abundance analysis. Peak annotation was performed using the ChipSeeker package v1.34.1^[55]^.

### LCMV Cl13 infection and restimulation of splenocytes

C57BL/6J mice were infected with LCMV Cl13 (2×10^6^ PFU/mouse). On day15 post infection, spleen was harvested, digested and single cell suspensions prepared using a mixture of collagenase/Dnase (Roche) prior to homogenation on a 100 mM filter using a butt-end of a syringe. RBCs were lysed for 2 minutes per spleen in 1× RBC lysis buffer. Following RBC lysis, B cells were depleted by magnetic bead separation using a CD19-positive selection II kit (EasySep). Splenocytes (2×10^5^ cells/well) were stimulated with 2 μg/mL LCMV-specific CD8 peptides (GP33-41, NP396-404 and GP276-286) and 5 μg/mL CD4 peptide (GP61-80) in the presence of Taz (0.125, 0.25, 0.5, 1.0 μM) or vehicle for 5 days. Plates were placed at 37 °C+ 5% CO2 at a 20 angle to increase cell to cell contact. Cells were analyzed on day5.

### Retroviral vector production

The plasmid construct of MSGC-14g2a-28z was obtained from Prof. Crystal L Mackall lab, transformed into XL1-Blue competent cells (Agilent, Cat#200249), and prepared using NucleoBond^®^ Xtra Midi Plus EF (Cat#740422.10). HEK293 Phoenix Ampho Packaging Cells were plated into a 10-cm dish 24h prior to the transfection. The medium was changed to fresh medium (9.0 mL) when the cell density reached 50-70% of confluence. The cells were incubated in the new medium for 1-2h for transfection. For transfection, linear polyethylenimine (PEI, 25 kDa, PolySciences, 30 ug) suspended in Opti-MEM (500 uL, Gibco) was added into the plasmid encoding HA-28z CAR (10 ug) suspended in Opti-MEM (500 uL, Gibco), incubated for 30 min at room temperature in the dark, and drop wised into the dish of cells. 6-8h after transfection, the medium was replaced by fresh medium (10.0 mL), and supernatants were collected at 36h and 60h after transfection. The retroviral vector in supernatants was concentrated using Retro-X™ Concentrator (Takara) following the manufacturer’s instructions.

### HA-28z CAR-T cell production and expansion

Cryopreserved human PBMCs were thawed and activated the same day with Dynabeads™ Human T-Activator CD3/CD28 for T Cell Expansion and Activation (Gibco, Cat#11131D) at 1:1 beads:cell ratio in T cell medium containing 150 IU/mL of IL-2. T cells were transduced with HA-28z retroviral vector at 24-36h after the activation and maintained at 0.5×10^6^-1×10^6^ cells/mL in T cell medium with 150 IU/mL of IL-2 for 4-5 days. CAR-T cells were stained with anti-mouse IgG F(ab’)_2_ (FITC, Jackson ImmunoResearch, Cat#515-095-072) and sorted out for expansion with 150 IU/mL of IL-2 in the presence of Tazemetostat (1.0 uM) or vehicle.

### Intracellular cytokine production of human T cells and CAR-T cells

On day13 after the 1^st^ activation (day7 after the 2^nd^ activation) of human peripheral blood monocytes, T cells (1 million/mL, 100 uL/well) were restimulated with coated anti-CD3 antibody (1 ug/mL, 100 uL/well, 4 °C overnight, Clone: OKT3, Biolegend) and soluble anti-CD28 antibody (2 ug/mL, Clone: CD28.2, Biolegend) in the culture medium containing 1×Brefeldin A (Biolegend, Cat#420601) in a 96-well plate for 5-7h. after incubation, intracellular cytokine staining was performed according to the method for flow cytometry analysis using the following antibodies: CD8a, Clone: HIT8a, IL-2, Clone: MQ1-17H12, IFNγ- Clone: 4S.B3.

For CAR-T cells, intracellular cytokine production was performed on day20 after the activation by co-culturing with targeting cells Nalm6-GD2. Briefly, CAR-T cells (1×10^5^ cells/well) and Nalm6-GD2 (1×10^5^ cells/well) were plated in Nalm6-GD2 culture medium containing 1×Brefeldin A (Biolegend, Cat# 420601) for 5-7h. After incubation, intracellular cytokine staining was performed according to the method for flow cytometry analysis using the same antibodies as above.

### CAR-T cell killing assay

4×10^4^ GFP^+^ Nalm6-GD2 cells were co-cultured with 1×10^4^ CAR-T cells in 200 uL culture mediu, in a black 96-well flat bottom plate for 24h. Triplicate wells were plated for each group. The fluorescence of GFP was recorded by an Agilent BioTek Synergy H4 Hybrid Microplate Reader.

### Statistics

All figures are representative of at least three experiments unless otherwise noted. All graphs report mean ±standard deviation (SD) values of biological replicates. For the SPICE analysis of cytokines, statistical significance was calculated using Wilcoxon signed-rank test. Statistical significance of two-group comparisons was calculated using a Student’s *t*-test. Analysis of multi-group comparisons was performed using one-way ANOVA and Dunnett’s post-test. *P*<0.05 was considered significant and is designated with an asterisk in all figures. R version 4.2.2 was used for the statistical analysis of RNA-seq and ChIP-seq data.

## Notes

### Competing Interest Statement

The authors have declared no competing interest.

