## Supplemental Figure1-9 for "Transient EZH2 suppression by Tazemetostat during *in vitro* expansion maintains T cell stemness and improves adoptive T cell therapy"

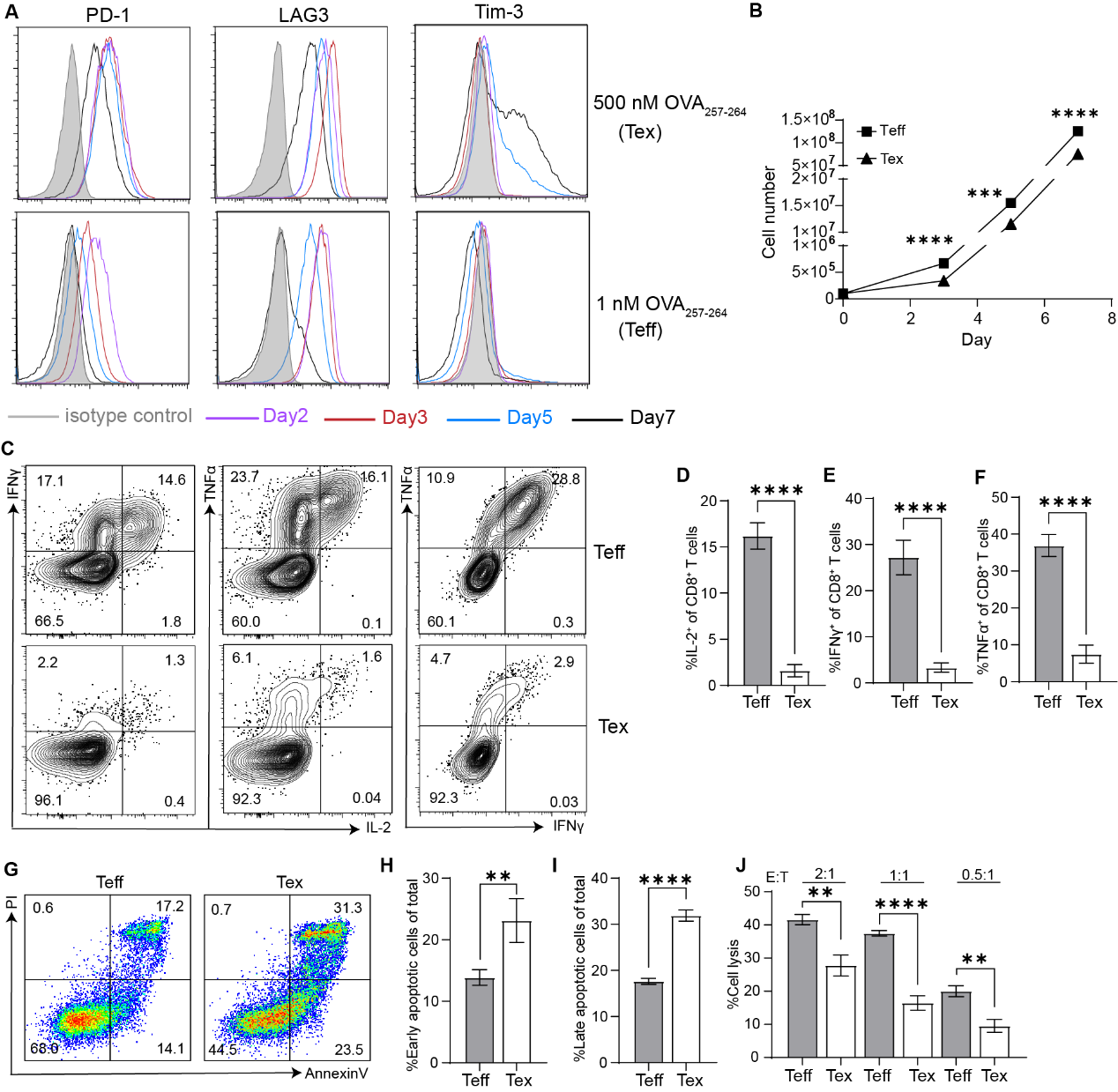

**Figure S1 Phenotype analysis of effector and exhausted OT-I T cells**. OT-I splenocytes were primed with 1 nM SIINFEKL peptide (OVA_257-264_) for 2 days and expanded for another 5 days in presence of 60 IU/mL IL-2, or 500 nM OVA_257-264_ for 3 days and expanded for another 4 days in presence of 60 IU/mL IL-2. Expression of inhibitory receptors from day 2 to day 7 (**A**). Proliferation curve of OT-I T cells (**B**). On day 7, OT-I T cells were re-stimulated with anti-CD3 (coated onto plate, 1 µg/mL for coating) and anti-CD28 (1 µg/mL in medium) antibodies and brefeldin A for 5-7h and stained with cytokine antibodies (**C-F**). Apoptosis analysis of OT-I cells re-stimulated with anti-CD3 (coated onto plate, 1 µg/mL for coating) and anti-CD28 (1 µg/mL in medium) antibodies for 24h (**G-I**). On day 7, cytotoxicity of OT-I T cells against 4MOSC1-SIINFEKL-GFP cells (**J**). ^ns^P > 0.05; *P < 0.05; **P < 0.01; ***P < 0.001; ****P < 0.0001; analyzed by student T-test. Tex: exhausted OT-I T cells, Teff: effector OT-I T cells.

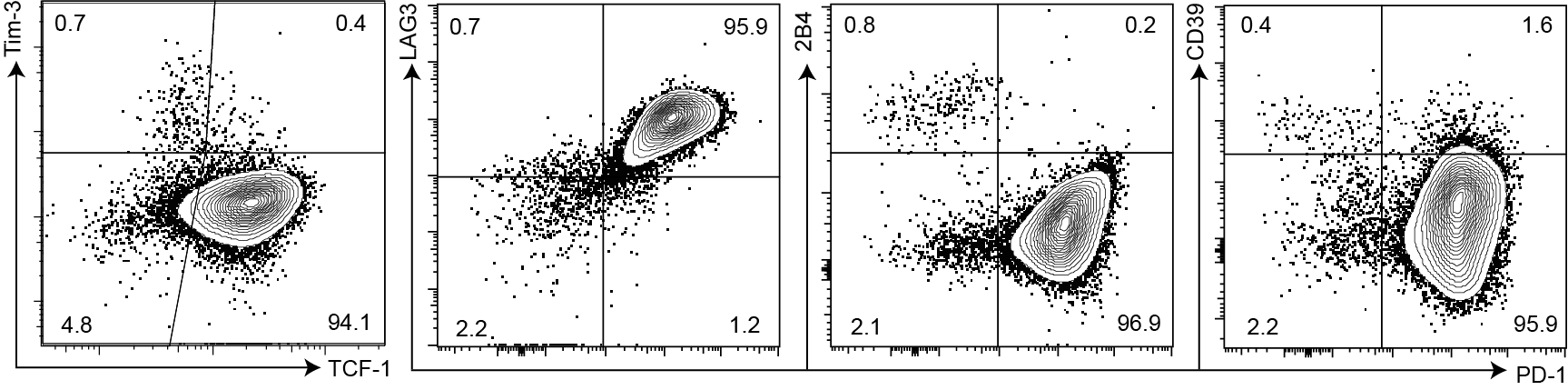

**Figure S2** OT-I splenocytes were stimulated with 500 nM OVA_257-264_ for 3 days in the presence of 60 IU/mL IL-2 and analyzed on day3.

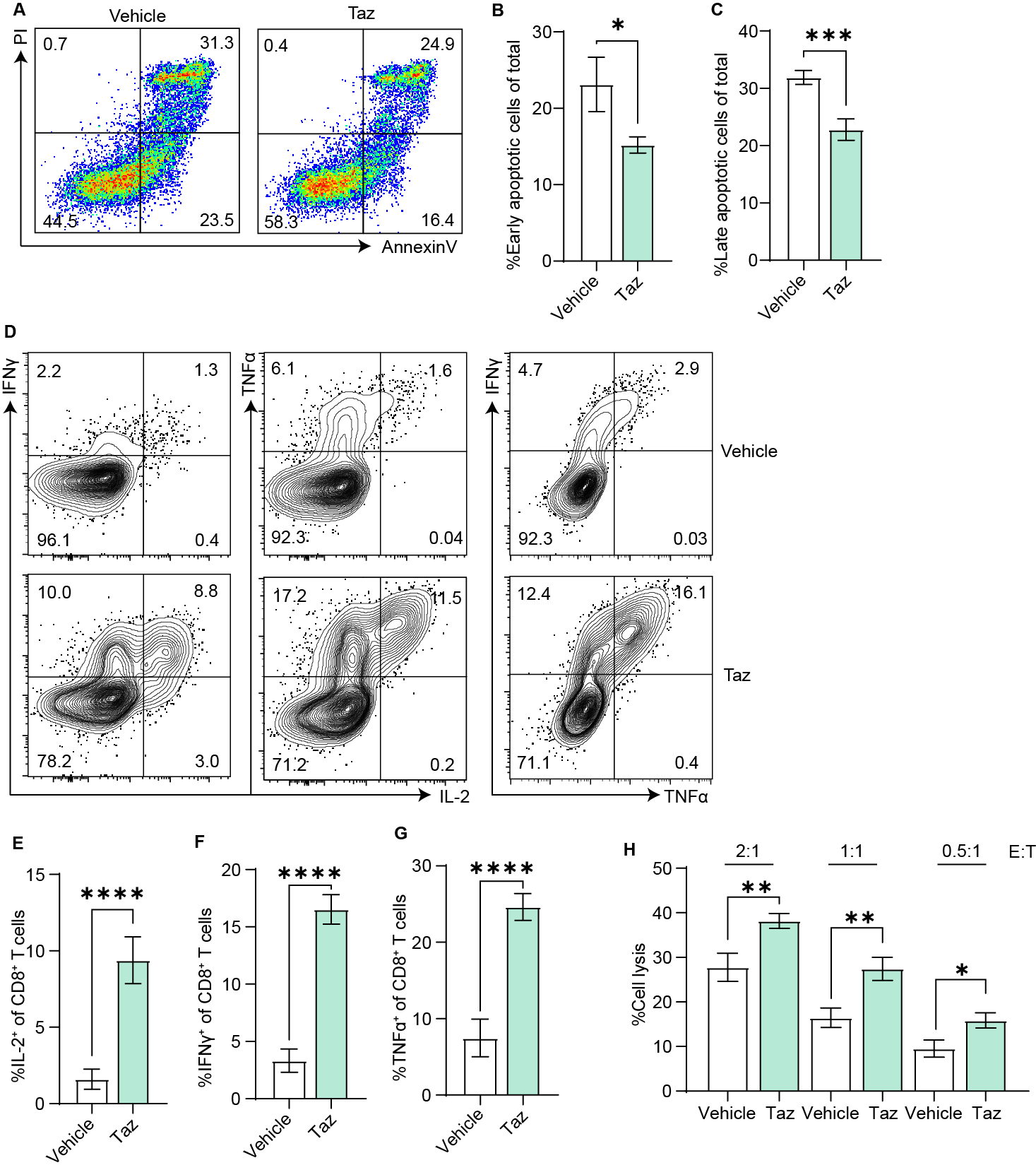

**Figure S3** Transient EZH2 inhibition by tazemetostat reduced the apoptosis (24 h) (A-C), increased cytokine production (D-G) of OT-I cells upon restimulation with anti-mouse CD3(coated on the plate)&anti-mouse CD28 (1 µg/mL in medium) and enhanced the cytotoxicity against targeting cells (20h) (H) on day 7. Three to four replicates for each group, ^ns^P > 0.05; *P < 0.05; **P < 0.01; ***P < 0.001; ****P < 0.0001; analyzed by student T-test.

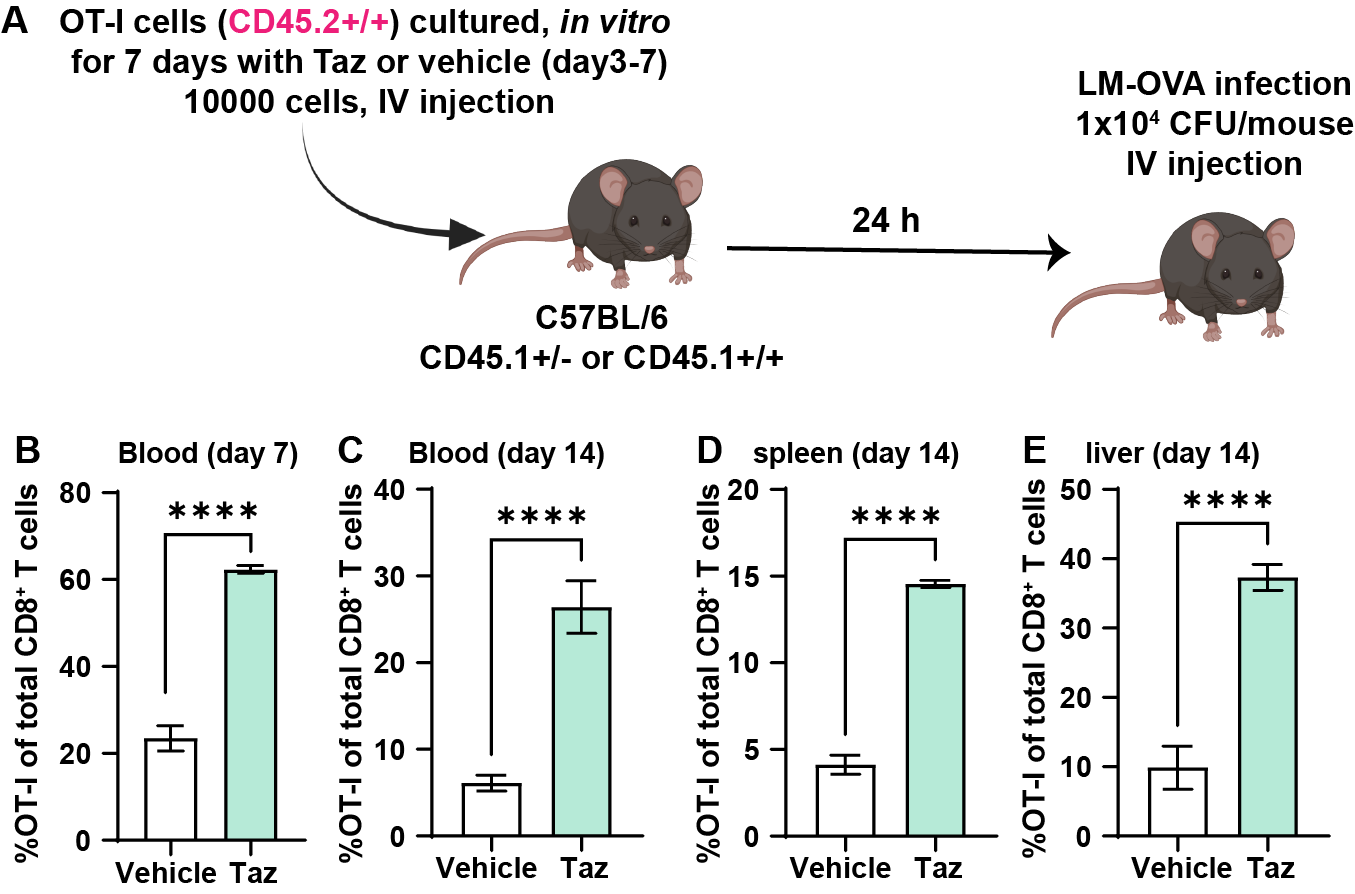

**Figure S4** **EZH2 inhibition by tazemetostat in OT-I cells during *in vitro* expansion enhanced *in vivo* response of OT-I cells upon LM-OVA infection.** OT-I splenocytes (CD45.2+/+) were activated with 500 nM OVA_257-264_ for 3 days and expanded in the presence or absence of Taz (1 uM) for 4 days (day3-day7). 24 h after OT-I CD8^+^ T cells (5000/mouse) were intravenously transferred into C57BL/6 mice (CD45.1+/+ or CD45.1+/-), mice were infected with 1×10^4^ CFU LM-OVA (A). Ratio of OT-I cells in CD8+ T cells in blood were analyzed on day 7 (B) and day 14 (C), in spleen (D) and liver (E) were analyzed on day 14. 4 mice for each group (n = 4), ^ns^P > 0.05; *P < 0.05; **P < 0.01; ***P < 0.001; ****P < 0.0001; analyzed by student T-test.

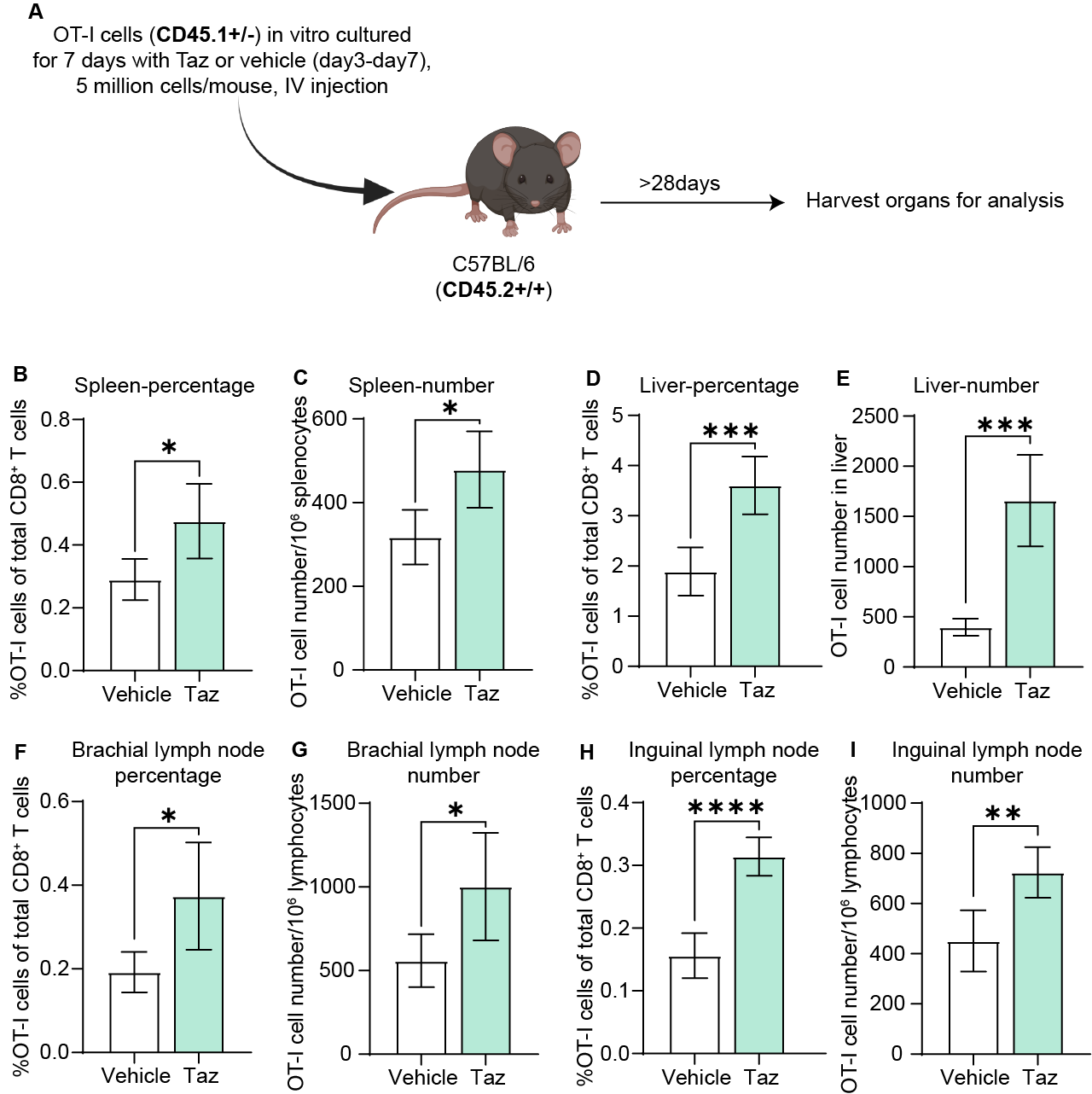

**Figure S5** **EZH2 inhibition by tazemetostat during *in vitro* expansion improved *in vivo* long-term survival of OT-I cells.** OT-I splenocytes (CD45.1+/-) were activated with 500 nM OVA_257-264_ for 3 days and expanded in the presence or absence of Taz (1 uM) for 4 days (day3-day7). 28 days after OT-I T cells (5 million cells/mouse) were intravenously transferred into C57BL/6 mice (CD45.2+/+), mice were sacrificed, and organs were harvested to quantify the percentage and number of OT-I cells. Percentages and number of OT-I cells in CD8^+^ T cells in spleen (B-C), liver (D-E), and lymph nodes (F-J) were analyzed. 5 mice for each group (n = 4), ^ns^P > 0.05; *P < 0.05; **P < 0.01; ***P < 0.001; ****P < 0.0001; analyzed by student T-test.

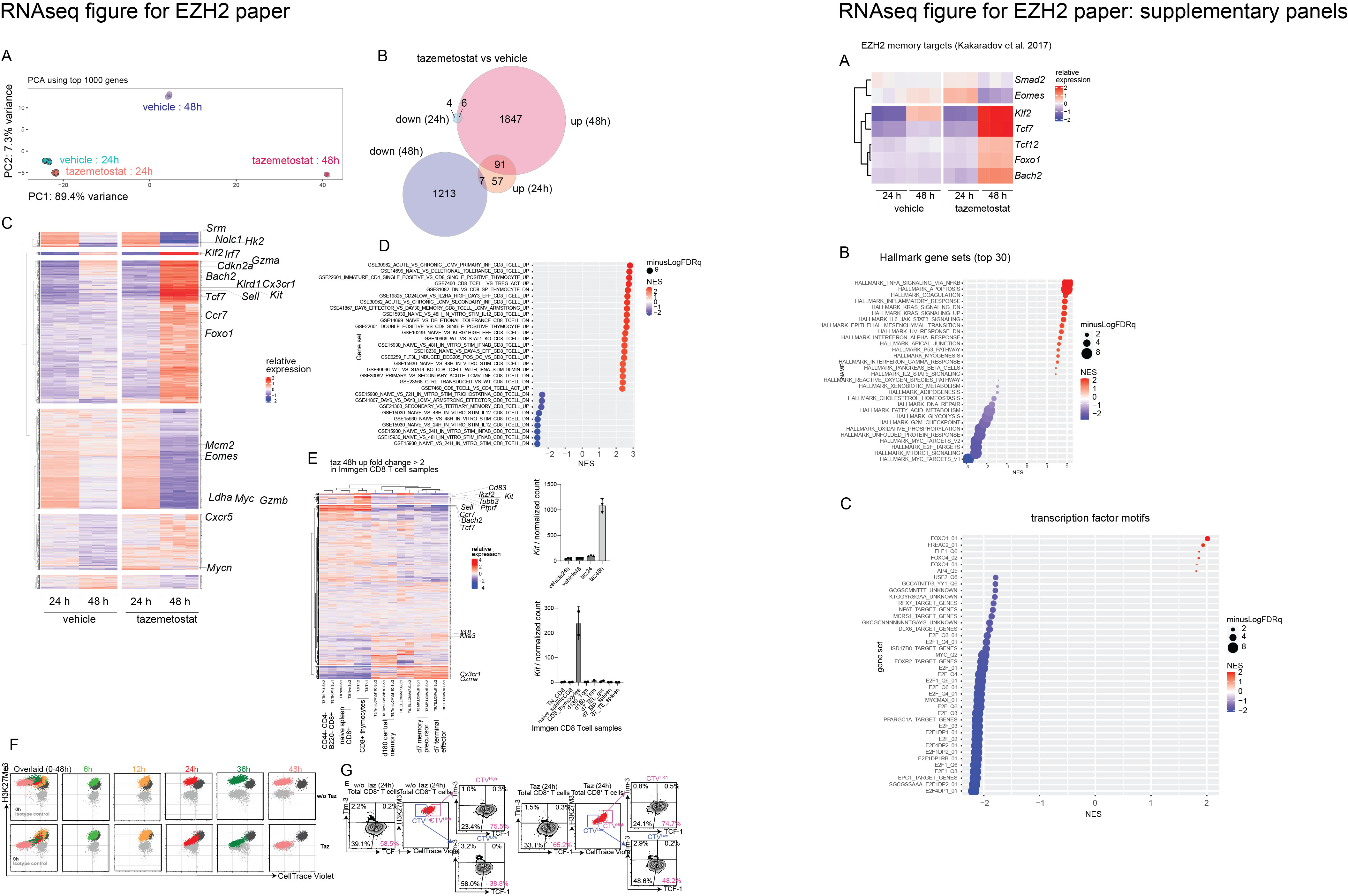

**Figure S6** OT-I splenocytes were stimulated with 500 nM OVA_257-264_ for three days and expanded in the presence of Taz or vehicle and RNA was isolated at 24h or 48h post treatment and analyzed by RNA-seq: (A) relative expression of known EZH2 targets in T cells based on Kakaradov et al^[^^1^^]^; (B-C) gene set enrichment analysis of Hallmark gene sets (B) and transcription factor motifs (C). Statistical normalization of RNA-seq reads was performed by DESeq2 (A); statistical comparison of experimental groups was performed by GSEA (B-C).

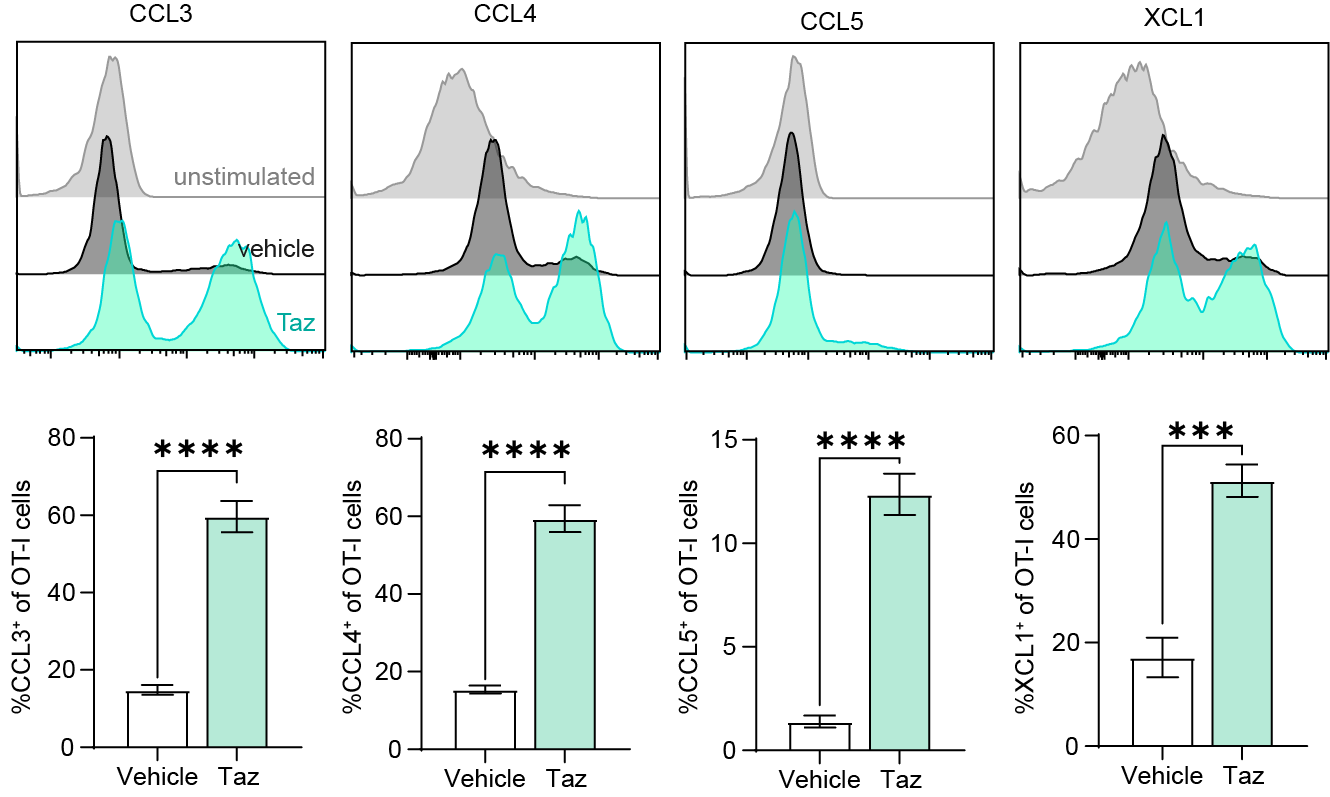

**Figure S7 Transient inhibition of EZH2 by tazemeostat increased chemokine production of OT-I cells.** OT-I splenocytes were activated with 500 nM OVA_257-264_ for 3 days and expanded in the presence or absence of Taz (1 uM) for 4 days (day3-day7). On day7, OT-I cells were re-stimulated with anti-CD3 (coated on the plate), anti-CD28 (1 µg/mL added in the medium) and brefeldin A for 5-7 h, and stained with anti-CCL3, anti-CCL4, anti-CCL5&anti-XCL1 antibodies.3 repeats for each group (n = 3), ^ns^P > 0.05; *P < 0.05; **P < 0.01; ***P < 0.001; ****P < 0.0001; analyzed by student T-test.

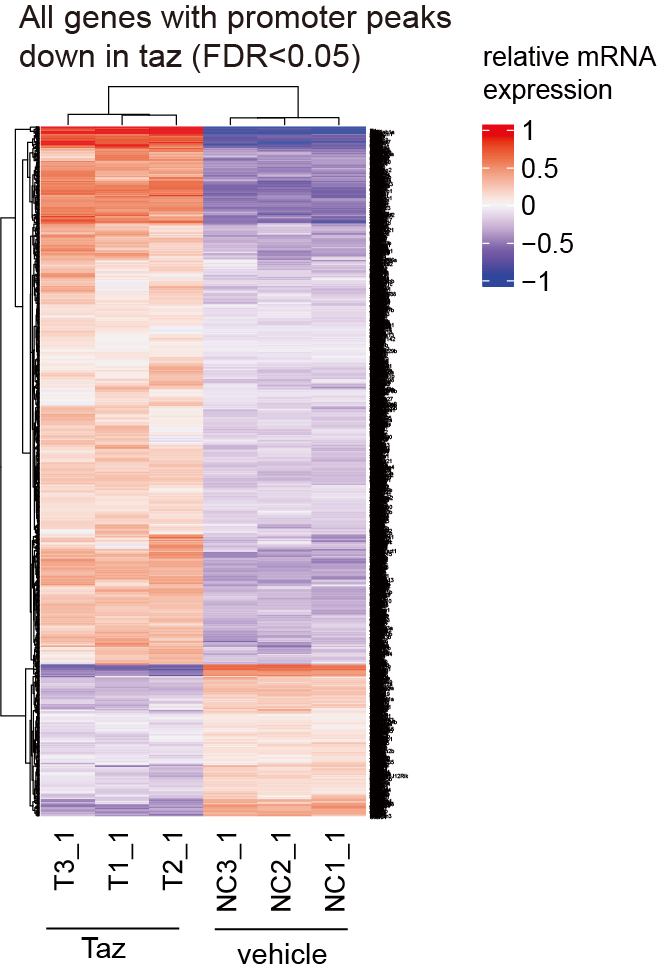

**Figure S8** Relative mRNA expression of genes whose promoter-associated peaks were reduced by Taz.
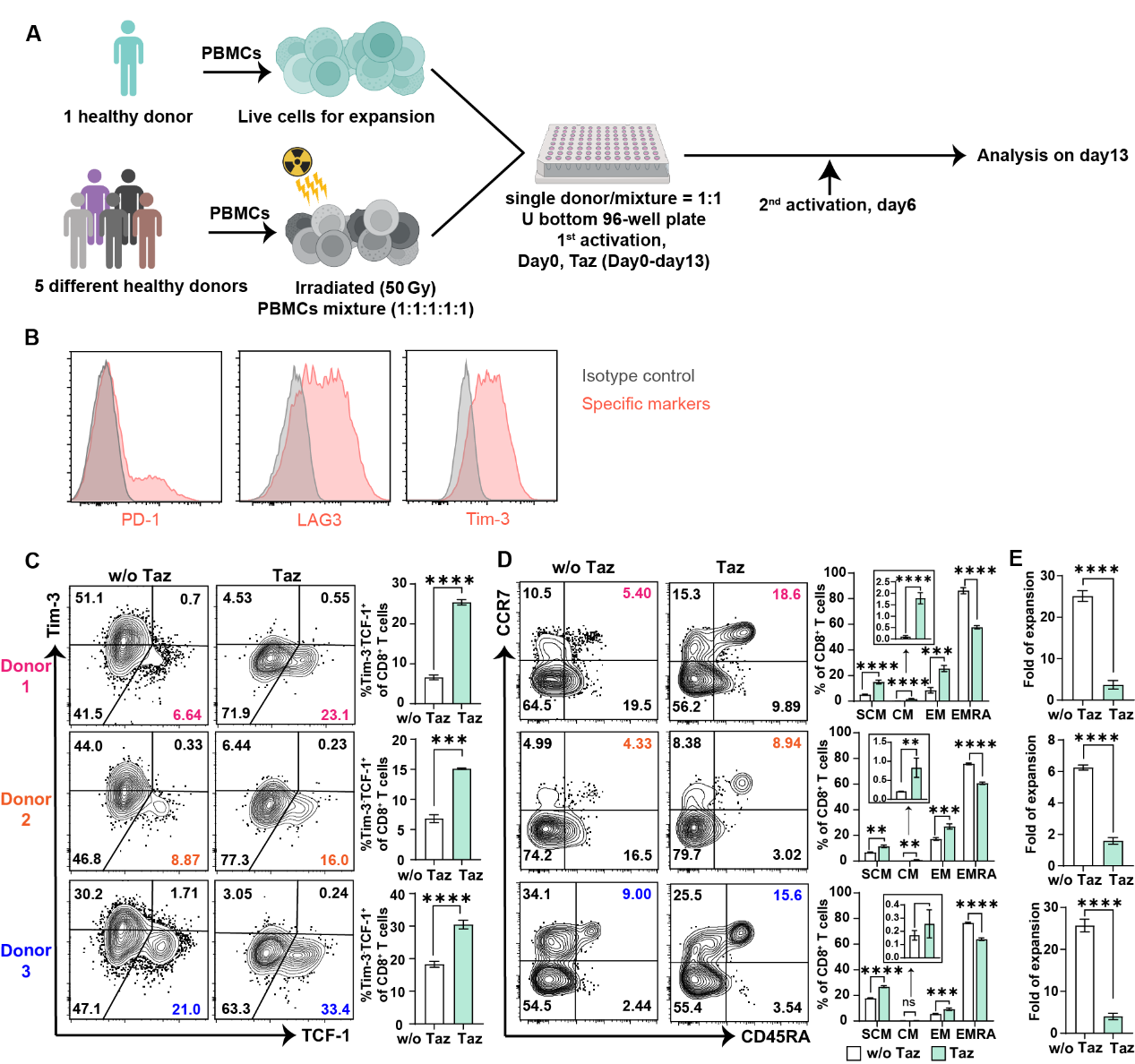

**Figure S9 Tazemetostat enhances desired properties in human PBMCs but inhibits the proliferation of hPBMCs**. Human PBMCs were activated with an irradiated (50 Gy) mixture hPBMCs from another five different donors on day0 and day6 and expanded with 300 IU/mL IL-2 in presence or absence of Taz (1uM, day0-day14) (A). Inhibitory receptors expression (B) of CD8^+^ T cells on day 13. Phenotype (C-D) analysis of CD8^+^ T cells and proliferation (E) analysis of total T cells in the presence of Taz or vehicle on day 13 for 3 donors. SCM: CD45RA^+^CCR7^+^, CM: CD45RA^-^CCR7^+^, EM: CD45RA^-^CCR7^-^, EMRA: CD45RA^+^CCR7^-^. 3 repeats for each group (n=3), nsP > 0.05; *P < 0.05; **P < 0.01; ***P < 0.001; ****P < 0.0001; analyzed by student T-test.
